# The chemotherapeutic drug methotrexate selects for antibiotic resistance

**DOI:** 10.1101/2020.11.12.378059

**Authors:** Jónína S. Gudmundsdóttir, Elizabeth G. A. Fredheim, Catharina I. M. Koumans, Joachim Hegstad, Po-Cheng Tang, Dan I. Andersson, Ørjan Samuelsen, Pål J. Johnsen

## Abstract

Understanding drivers of antibiotic resistance evolution is fundamental for designing optimal treatment strategies and interventions to reduce the spread of antibiotic resistance. Various cytotoxic drugs used in cancer chemotherapy have antibacterial properties, but how bacterial populations are affected by these selective pressures is unknown. Here we test the hypothesis that the widely used cytotoxic drug methotrexate affects the evolution and selection of antibiotic resistance through the same mechanisms as the antibiotic trimethoprim. We show that methotrexate can select for trimethoprim resistance determinants located on the chromosome or a plasmid in clinical strains of *Escherichia coli*. Additionally, methotrexate can co-select for virtually any antibiotic resistance determinant when present together with trimethoprim resistance on a multidrug-resistance clinical plasmid. These selective effects occur at concentrations 40- to >320-fold below the methotrexate minimal inhibitory concentration for *E. coli*, suggesting a selective role of methotrexate chemotherapy for antibiotic resistance in patients that strongly depend on effective antibiotic treatment.

**Significance statement:** The presented data show that methotrexate has the potential to select for virtually any given antibiotic resistance gene when genetically linked to trimethoprim resistance. This study highlights the need for increased awareness of the presence of acquired antibiotic resistance determinants in the gut of patients with impaired immunity undergoing methotrexate treatment to preserve the effects of downstream antibiotic treatments.

## Introduction

Antibiotic resistance is a major risk factor for patients with impaired immunity, such as cancer patients, and often a patient’s survival depends on antibiotic treatment to reduce the risk for hospital-acquired infections during chemotherapy^1,2^. Several cytotoxic drugs used in cancer chemotherapy are known to both elevate bacterial mutation rates and have direct antimicrobial properties^3,4^. It has been proposed that cancer chemotherapy may drive *de novo* antibiotic resistance evolution through SOS induced mutagenesis^5^, and some reports have provided support for this hypothesis^6,7^. Recently, the effects of non-antibacterial drugs on bacteria typically found in the human gut were thoroughly explored and cytotoxic drugs were reported to cause the most severe alterations of the microbiota^8^. Taken together, these studies suggest that cytotoxic drugs affect survival of human gut commensals, they may increase the evolvability of bacterial populations, and lead to reduced susceptibility towards drugs used to treat cancer in bacteria. How bacterial populations respond to selective and co-selective pressures exerted by individual cytotoxic drugs and the implications for antibiotic resistance selection and spread is unknown. Thus, there is an urgent need to understand these potential collateral effects of cancer chemotherapy to ensure effective antibiotic treatment for a large group of immunocompromised patients. Moreover, cytotoxic drugs may constitute a previously unrecognized target for intervention to limit the selection and spread of antibiotic resistance.

Methotrexate (MTX) is widely used in treatments including but not limited to; cancer of the breast, skin, head, neck, and lung as well as many inflammatory diseases, such as rheumatoid arthritis^9^. We specifically targeted resistance towards trimethoprim (TMP), since both drugs act through inhibiting the dihydrofolate reductase enzyme in bacteria and eukaryotic cells, central in DNA synthesis^10^. TMP in combination with sulfamethoxazole is among the most frequently used antibiotics in the treatment of urinary tract infections and is recommended as first line treatment internationally^11^. Our target organism in this study is *Escherichia coli*, the most common agent of nosocomial infections world-wide^12^. *E. coli* is known to display intrinsic resistance towards methotrexate through AcrAB-TolC mediated efflux^13^, however TMP is not a substrate for this efflux system.

Previous studies have focused on the abilities of MTX and other non-antibiotics to inhibit bacterial growth^8^. These approaches have provided valuable insights on the effects of non-antibiotics as modulators of the intestinal flora but lacked the necessary resolution to detect more subtle selective effects on acquired antibiotic resistance determinants.

Here, we hypothesize that despite the demonstrated *E. coli* intrinsic MTX resistance^13^, MTX can affect antibiotic resistance evolution in *E. coli*, due to the shared molecular target with TMP. We show that MTX selects for acquired bacterial TMP resistance (TMP^R^) and co-selects for other antibiotic resistance determinants when co-residing on a mobile genetic element. Exposure to a wide concentration range of MTX selects for mutations identical to those emerging during TMP selection in clinical isolates of *E. coli*. Moreover, we show that the minimum selective concentrations (MSCs) of MTX and positive selection for chromosomal and plasmid-mediated TMP^R^ determinants occurs at concentrations 40- and >320-fold below the MTX minimum inhibitory concentrations (MICs), respectively.

## Methods

### Bacterial strains and growth conditions

Bacterial strains used in this study and their assigned reference numbers are listed in Table S1. Most strain constructs were derived from the *E. coli* K-12 strain MG1655 (DA4201/DA5438), or for cloning purposes from *E. coli* DH5α (MP18-09). Two isogenic pairs of clinical *E. coli* strains, where one strain is trimethoprim susceptible (TMP^S^) and the other is TMP^R^, were used for fitness measurements. In the first pair, TMP^R^ in *E. coli* K56-2 (MP06-01) is caused by one intragenic point mutation T>A (W30R) in the *folA* gene, and one in its promotor region (P_*folA*_, C>T 58 base pairs (bp) upstream of the gene)^14,15^ (MP06-05). In the second pair, *E. coli* K56-75 (MP06-41) harbouring the pG06-VIM-1 multi-drug resistance plasmid represents the TMP^R^ counterpart (MP05-31). This plasmid encodes, amongst seven other antibiotic resistance genes, two different *dfrA* genes (*dfrA1* and *dfrA12*) as putative TMP^R^ determinants as well as a carbapenem resistance determinant (*bla*_VIM-1_)^16^. For accurate MSC measurements we constructed pairs of fluorescently tagged (*yfp* and *bfp*) *E. coli* MG1655 strains containing the same resistance markers as described above. For details concerning growth conditions, susceptibility testing and strain constructions see protocols in supplementary information (SI).

### Fitness measurements

Growth rates were determined using a BioscreenC MBR reader (Oy Growth Curves Ab, Ltd). The growth rate calculations were done using the Bioscreen Analysis Tool BAT 2.1^17^ (see SI for detailed protocol). Competition experiments were performed in six biological replicates, including dye swaps, using the fluorescently tagged strain pairs described above, containing *folA* or pG06-VIM-1 mediated TMP^R^. Both competition experiments, MSC calculations and competitions performed at different starting ratios were done as previously described in^18^ (see SI for detailed protocols).

### Experimental evolution

To examine the effect of MTX presence on TMP^R^ evolution, strain K56-2 (MP06-01) was serially passaged in liquid cultures with MTX supplemented at a concentration slightly above the estimated MSC. Initially, ten independent overnight cultures were started from independent colonies on separate agar plates. From the original overnight cultures, ~10^3^ cells were used to start ten independent lineages in 1 mL batch cultures containing 400 μg/mL MTX (lineages 1-10). Every 12 h for 25 days, the lineages were serially passaged by 1,000-fold dilution, allowing for ~500 generations of growth. Every ~50 generations the populations were frozen down at −80°C for downstream analysis. For detailed protocols concerning experimental evolution, selective plating on high concentrations of MTX and whole genome sequencing see SI.

## Results

### Methotrexate selects for pre-existing trimethoprim resistance determinants

We determined the MICs of MTX in clinical and laboratory strains of *E. coli* (Table S1). Initial experiments revealed variable, but high MTX MICs (Table S2) consistent with previous reports demonstrating that *E. coli* displays intrinsic resistance towards MTX due to AcrAB-TolC mediated efflux^8,13^. We also observed that the MTX MIC was dependent on the presence of TMP^R^ determinants. All isolates with a functional TMP resistance determinant and increased TMP MIC showed consistently higher MTX MICs (>32 mg/mL) than TMP susceptible isolates (4-32 mg/mL), indicating possible co-selective abilities of the two drugs.

Antibiotic resistance selection and co-selection have traditionally been assumed to occur between the MICs of susceptible and resistant isolates within a bacterial population (known as the selective window)^18^. However, several reports unequivocally show that antibiotic resistance selection and co-selection can occur at concentrations several hundredfold below the MIC of a susceptible isolate (known as sub-MIC)^18–20^. To test how sub-MICs of MTX affect bacterial fitness, we measured exponential growth rates for two pairs of clinical isogenic TMP^R^ and TMP^S^ *E. coli* across a wide MTX concentration span. One pair with TMP^R^ located on the chromosome (two mutations associated with *folA*)^14^ and one pair with TMP^R^ (*dfrA*) located on the MDR plasmid pG06-VIM-1^16^. TMP^S^ strains displayed sharply declining growth rates between 1 and 2 mg/mL of MTX, whereas the TMP^R^ strains remained unaffected (Figure S1). These results suggest a selective benefit during MTX exposure for TMP^R^ strains at concentrations below the observed MTX MIC of the TMP^S^ clinical isolates.

The MSC describes pharmacodynamically the lowest concentration where selection for resistance occurs^18^. To determine the MSC for MTX, we constructed a fluorescently tagged pair of *E. coli* MG1655 strains to enable accurate separation between the two in mixed populations. In these backgrounds, we introduced TMP^R^, either through SNPs (*folA*) using genome engineering or through the pG06-VIM-1 plasmid. The isogenic TMP^R^ and TMP^S^ strain pairs were competed head-to-head by serial passage for 30 generations and the ratio of TMP^R^:TMP^S^ was determined over time using flow cytometry. From this data the MSC was estimated (Tables S3-S5)^18^. *E. coli* with chromosomal *folA* mutations had an MSC of 200 μg/mL (1/40 of the MIC of MTX) whereas those harboring the plasmid were close to the detection limit of the assay^18^ and we conservatively estimated the MSC to be <25 μg/mL (less than 1/320 of the MIC of MTX) (Figure 1). Taken together, our data strongly suggest that selection for TMP^R^ occurs at MTX concentrations far below the estimated MTX MIC.

**Figure 1:**
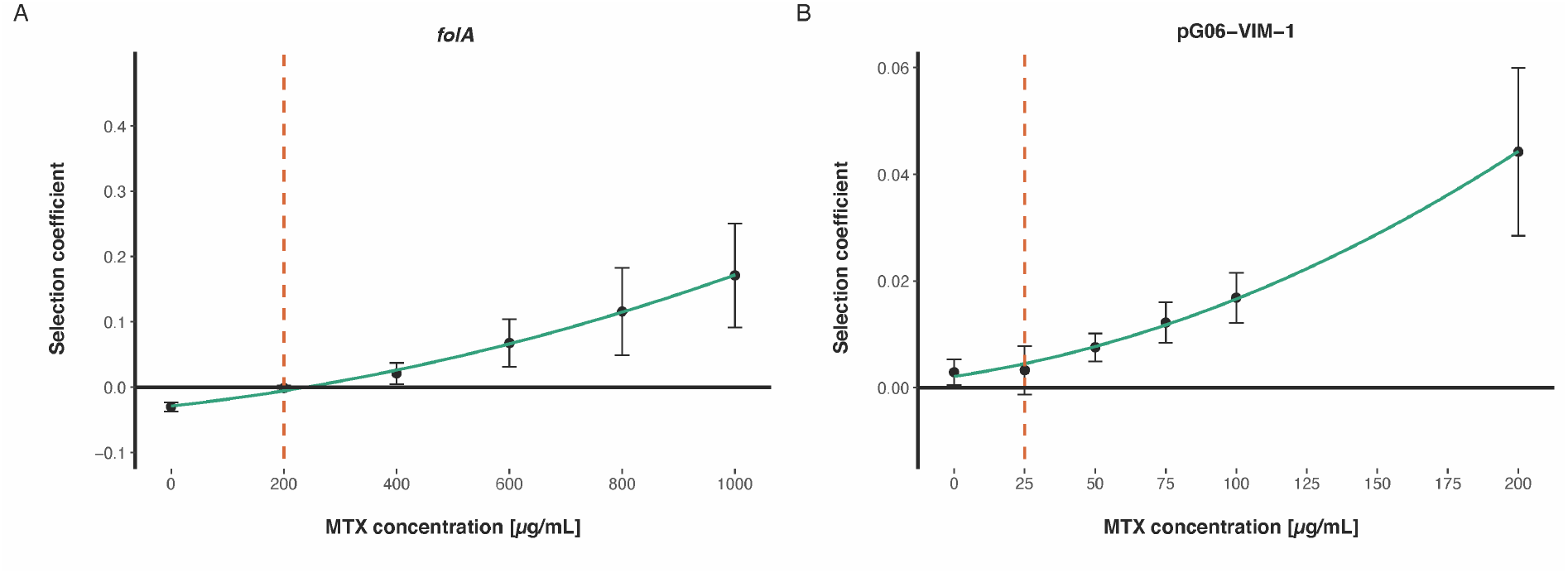
Selection coefficients as functions of MTX concentrations from competition experiments between TMP^R^ and TMP^S^ isogenic strains. The MSC is defined as the concentration where the selection coefficient equals zero. The MSC of *E. coli* MG1655 harboring **(A)** two chromosomal *folA* mutations (MP18-04 and MP18-07) is set to 200 μg/mL, and **(B)** the MDR pG06-VIM-1 plasmid encoding *dfrA12* (MP18-05 and MP18-08) is conservatively set at 25 μg/mL. Dashed lines represent the set MSC, bullets the average selection coefficients and error bars the standard deviations.

### Sub-MICs of methotrexate promotes invasion of trimethoprim resistance determinants even when rare in *E. coli* populations

Exploring MTX-selective dynamics further, we asked if TMP^R^ determinants could invade the population at lower initial densities to exclude potential bias from the 1:1 ratio in the competition experiments. We started competition experiments from frequencies as low as 10^−4^ of the TMP^R^ strains, at concentrations slightly above the estimated MSC of MTX (400 μg/mL for *folA* mediated resistance and 75 μg/mL for pG06-VIM-1 mediated resistance) and followed the change in ratios over 30 generations of growth (Figure 2). Both chromosomal and plasmid mediated TMP^R^ determinants were able to invade, even when initially rare in their respective populations, strongly suggesting that the MTX selective effects are independent on initial frequencies of resistant and susceptible strains during competition experiments.

**Figure 2:**
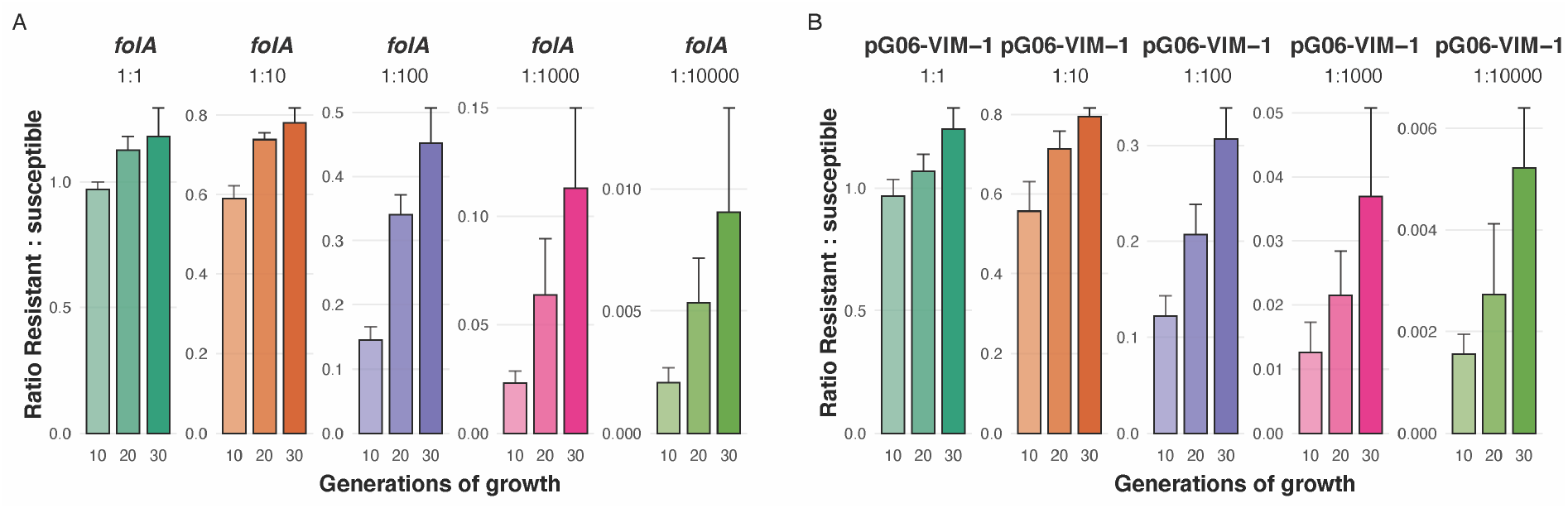
Competition experiments during sub-MIC MXT exposure at different initial frequencies of TMP^R^ strains. The change in TMP^R^:TMP^S^ ratios over 30 generations of growth at MTX concentrations slightly above the MSCs where *E. coli* MG1655 harbors **(A)** two *folA* mutations (MP18-07), and **(B)** the pG06-VIM-1 plasmid (MP18-08) were competed against a differently tagged, isogenic susceptible strain (DA56507) at 1:1, 1:10^−2^, 1:10^−3^ and 1:10^−4^ starting ratios. Error bars represent the standard deviation of the average ratio.

### Methotrexate co-selects for resistance determinants on a multi-drug resistance plasmid

The *dfr*-genes represent a common TMP^R^ mechanism in *E. coli* and these genes are frequently located on mobile genetic elements such as integrons and plasmids. Given that MTX selects for *dfr*-mediated TMP^R^, co-selection of other genetically linked resistance genes is likely. To show this, we used the MDR pG06-VIM-1 plasmid harboring *dfrA1* and *dfrA12* along with multiple resistance determinants including four aminoglycoside resistance genes and the *bla*_VIM-1_ carbapenemase gene conferring resistance to broad-spectrum β-lactams including carbapenems^16^. To assess the stability of pG06-VIM-1 in our strains competing in the presence of MTX, *E. coli* K56-75 harboring the plasmid (MP05-31) was serially passaged in batch cultures with 400 μg/mL MTX supplemented for 50 generations. The lineages were then plated on non-selective agar and 100 colonies from each lineage tested for reduced susceptibility towards ampicillin, TMP, streptomycin and spectinomycin. The results revealed complete phenotypic stability across all three lineages, confirming MTX mediated co-selection of plasmid mediated MDR.

To verify that the TMP^R^ determinants on the MDR pG06-VIM-1 plasmid is the primary mediators of MTX resistance and selection (Table S2), both *dfrA1* and *dfrA12* were isolated from the plasmid and cloned onto an expression vector and the effects of the individual genes measured. Of the two genes, only *dfrA12* was shown to give the same resistance pattern for TMP as well as MTX as the pG06-VIM-1 plasmid (Figure 3), and the lack of detectable phenotype for *dfrA1* (MP18-11) is likely due to a start codon frameshift mutation^16^.

**Figure 3:**
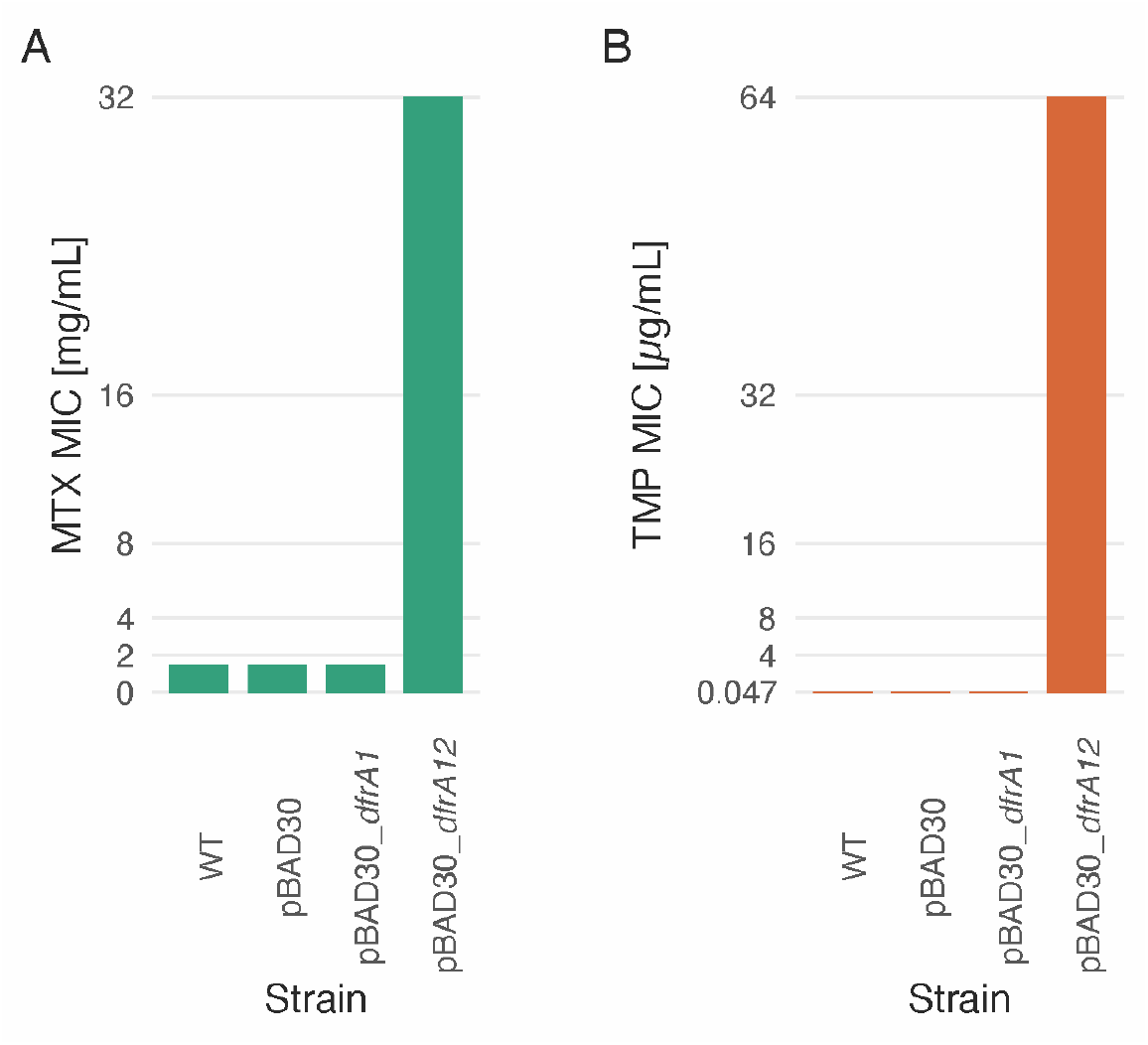
MTX and TMP MIC of *E. coli* DH5α expressing *dfrA1* or *dfrA12.* The MTX **(A)** and TMP **(B)** MIC for the wild type (WT) *E. coli* DH5α (MP18-09) compared to the strain harboring the empty pBAD30 (MP18-10) expression vector as well as strains with pBAD30 with different *dfrA* genes expressed under the inducible expression control by the pBAD promotor (MP18-11 and MP18-12). The detection limit of the assay is 64 μg/mL for TMP and 32 mg/mL for MTX. For both drugs, the MIC of *E. coli* DH5α expressing *dfrA*12 (MP18-12) exceeded the detection limit whereas the strain expressing *dfrA1* (MP18-11) has the same MICs as the WT strain. Showing that MTX and TMP resistance conferred by the pG06-VIM-1 plasmid is caused by the *dfrA12* gene.

### Methotrexate selects for *de novo* trimethoprim resistance

We further examined whether exposure to MTX could lead to *de novo* TMP^R^ evolution. We selected spontaneous mutants from overnight cultures with and without exposure to MTX, plated on selective agar at high MTX concentrations and tested for TMP cross-resistance (Figure S2, Table S6). *E. coli* K56-2 isolated at 16 and 32 mg/mL MTX (MP18-17 to MP18-28) displayed increased MICs of TMP close to or above the clinical breakpoint^21^, clearly demonstrating selection for TMP^R^ by MTX. DNA sequencing of the resistant isolates revealed two different mutations in the *folA* promoter, previously reported to result in TMP^R22^, as well as a single mutation in the *marR* gene (Table S6-S11).

Finally, we asked if exposure to sub-MICs of MTX close to the estimated MSCs would select for *de novo* TMP^R^ mutations in a susceptible *E. coli* population. Starting from 1,000 cells to minimize the probability of pre-existing mutants, we grew ten independent lineages of the *E. coli* K56-2 strain at 400 μg/mL MTX for 500 generations. The frequency of TMP^R^ was determined every 50 generations. TMP^R^ ascended in frequency in 2/10 lineages at different rates and time-points during the first 250 generations before they were outcompeted by a different set of mutants with reduced susceptibility to MTX and no cross-resistance to TMP (Figure 4, Table S12). These experiments show that MTX exposure can select for *de novo* TMP^R^, both at high and sub-MIC concentrations.

**Figure 4:**
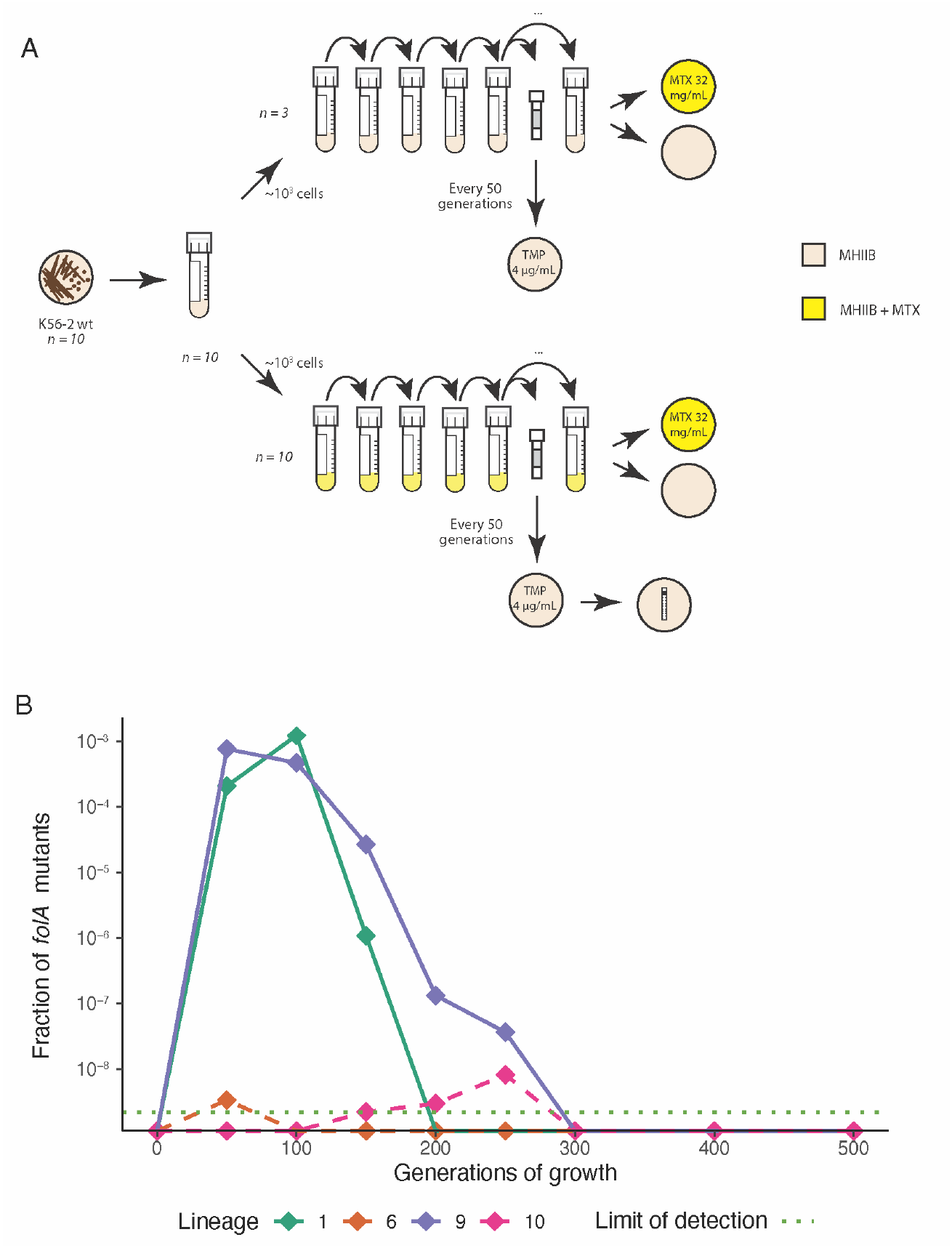
Evolution of TMP^R^ during MTX exposure for 500 generations. **(A)** Sub-MIC evolution experimental set-up. Ten biological replicates of K56-2 (MP06-01) were evolved for ~500 generations with 400 μg/mL MTX and three biological replicates without drug. All lineages were screened for TMP^R^ every 50 generations. After 500 generations all end-point populations were plated on 32 mg/mL MTX. All populations were able to grow at 32 mg/mL MTX, but not a single clone isolated conferred TMP^R^, strongly suggesting that reduced susceptibility to MTX with no cross-resistance to TMP evolved in the endpoint populations. **(B)** Fractions of TMP^R^ *folA* mutants isolated every 50 generations from the lineages where these were detected. The detection limit of the assay was ~2×10^−9^. Solid lines represent the two lineages where TMP^R^ emerged and ascended in frequency whereas dotted lines indicate spontaneous mutants.

### Pharmacokinetic approximations

To assess pharmacokinetic relevance, we attempted to estimate the MTX concentration range likely to be found in the intestine of patients undergoing MTX treatment. Limited information is available on gut MTX concentrations following intravenous administration during cancer treatment, as pointed out by others^8^. Pharmacokinetic data reveal that up to 90% of administered MTX is renally excreted^23^ and we assume that the remaining ~10% of the dose constitutes the upper limit of the concentration range present in the human intestine. The lower limit is set to 2% of the dose based on the mean ^3^H labelled MTX concentrations measured in stool samples from nine patients receiving MTX intravenously^24^. From this, we set a 24 hour transition time in a total volume of 0.6 L ^8^ and calculated the dose (*d*) required to achieve MSC in the human intestine from:

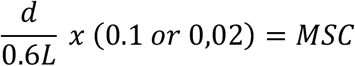

Estimated doses needed to reach intestinal MSCs assuming 2% and 10% fecal MTX concentrations were from 0.15g to 0.75g for plasmid-mediated TMP^R^ and from 1.2g to 6g for chromosomal *folA* mutations. Thus, assuming close to 2m^2^ body surface in grown-up patients^25^ estimates of MSC for plasmid-mediated TMP^R^ translates to dosing regimens from 75-375 mg/m^2^ and from 0.6-3 g/m^2^ for the chromosomal *folA* mutations. These approximations indicate that our MSC estimates are relevant for patients receiving high dose MTX treatment (1-12 g/m^2^)^26^. A recent study used a different approach estimating gut MTX concentrations following oral administration during treatment of rheumatoid arthritis^27^. Their data suggested MTX concentrations as high as 100 μg/mL are found in the gut, suggesting that our estimated MSC for plasmid-mediated antibiotic resistance determinants (25 μg/mL) is well within this concentration range.

## Discussion

Here we show that exposure to the cytotoxic drug MTX affects selection and evolution of TMP^R^ determinants at clinically relevant concentrations. Notably, MTX can mediate selection of any antibiotic resistance determinant when TMP^R^ is co-localized on a mobile genetic element across a wide concentration gradient. Arguably, this potentially important side-effect of MTX treatment has been previously unrecognized, as studies on the effects of non-antibiotic drugs, including MTX, have either focused on bacterial growth inhibition or used drug concentrations around the MIC^8,28–30^, with a few exceptions^19,31^. Using the approaches outlined here, including high resolution mixed culture competition experiments, allow determination of the true MTX selective window ranging from the MSC to the MIC^18^. This is particularly relevant for non-antibiotics for which bacteria display reduced susceptibility. In *E. coli*, MTX is a substrate for the AcrAB-TolC efflux pump^13^ and selective effects as those demonstrated here would not have been detected in classical susceptibility and/or growth assays in bacterial monocultures. This was recently supported in an *E. coli* chemical genetic screen where clear growth inhibitory effects of MTX, as well as for a range of other non-antibiotics, were only demonstrated in a *tolC* knock-out mutant (i.e. in a mutant lacking the intrinsic mechanism of resistance)^8^.

Given that many cytotoxic drugs are structurally similar to antibiotics (e.g. doxorubicin/tetracyclines), or target similar key processes as the major antibiotic groups (e.g. DNA/protein synthesis) it is possible that cancer chemotherapy may lead to increased levels of antibiotic resistance in a vulnerable patient group that very often rely on efficient antibiotic treatment for survival. To acquire a deeper understanding of the evolutionary potential of novel, non-antibiotic drivers of antibiotic resistance the approaches presented here are essential. These approaches need to be combined with an improved understanding of the intestinal pharmacokinetics of MTX and other cytotoxic drugs, possible effects of co-administered drugs such as leucovorin mediated MTX rescue^32^, and their interactions with the human microbiome. Such knowledge could allow identification of antibiotic + non-antibiotic drug combinations that should be avoided to preempt resistance evolution.

Recent studies have showed that non-antibiotics can increase mutation rates^6,7^ and promote horizontal gene transfer^31,33^. Taken together with the data presented here it is clear that non-antibiotic drugs have the potential to affect the evolution, selection, and spread of antibiotic resistance determinants. It is also clear that without a better understanding of these potential drivers of antibiotic resistance, interventions including those targeting human consumption of antibiotics may be negatively affected.

## Acknowledgments

We thank: Fredrik Sund, MD at The University Hospital North Norway for valuable input on medical aspects of MTX. Ass. Prof. Daniel Rozen, Leiden University for valuable discussions. Prof. Natasa Skalko-Basnet and Ass. Prof. Ann Mari Holsæter, UiT The Arctic University of Norway, for training in handling of cytostatic drugs. Funding: PJJ was supported by a joint grant from UiT The Arctic University of Norway and the Northern Norway Regional Health Authority (A23270). DIA was supported by the Swedish Research Council (grant 2017-01527).

## Author contributions

The study was designed by JSG, EGAF, ØS and PJJ. Strain constructions were done by PT and JSG. Experiments were conducted by JSG, EGAF and CIMK. WGS analysis was done using a bioinformatic pipeline designed by JH. WGS data was verified by JH and JSG. Experimental data was verified by JSG and PJJ. Formal data analysis and data visualization was done by JSG. The first draft of the manuscript was written by JSG and revised by PJJ. All authors revised subsequent version and provided key edits to the manuscript. Funding acquisition and resources were provided by PJJ and DIA.

## Data availability

WGS data are available at NCBI (BioProject PRJNA677979).

