## Supplementary Information for "The chemotherapeutic drug methotrexate selects for antibiotic resistance"

**Supplementary Figures**

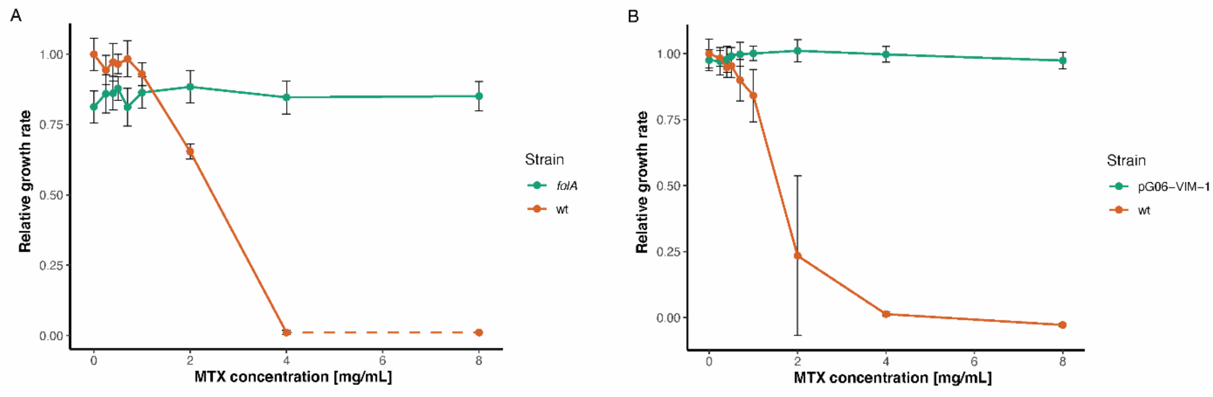


**Figure S1: Relative growth rates of TMP$^{R}$ strains compared to their susceptible WTs as functions of MTX concentration. (A)** Chromosomally mediated TMP$^{R}$ through *folA* mutations allows the K56-2 (MP06-05) strain to remain unaffected at MTX concentrations as high as 8 mg/mL whereas the TMP susceptible WT strain (MP06-01) shows a steep decrease in relative growth rates already between 1 and 2 mg/mL MTX. As we are close to the MIC of MTX for the TMP$^{S}$ strain at 8 mg/mL MTX, growth rate measurements were difficult to interpret due to noise in the measurements. Dotted line indicate that the value is set close to zero. **(B)** Plasmid-mediated TMP$^{R}$ through *dfrA* located on pG06-VIM-1 (MP05-31) allows the K56-75 strain to remain unaffected at MTX concentrations as high as 8 mg/mL whereas the TMP susceptible WT strain (MP06-41) shows a steep decrease in relative growth rates already at 0.7 mg/mL MTX. The error bars represent the standard deviation of the mean growth rate of 10 replicates (5 biological replicates with 2 technical replicates each).

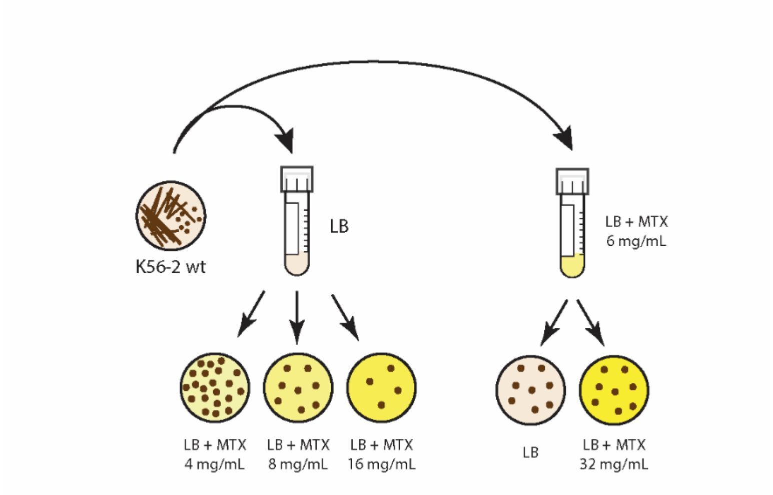


**Figure S2: Schematic of the experimental set-up used during selective plating at high MTX concentrations.** The MTX susceptible K56-2 strain (MP06-01) was grown overnight in LB without any drug present before plating on increasing concentrations of MTX, 4, 8 and 16 mg/mL. In a similar manner the K56-2 strain (MP06-01) was grown in LB + 6 mg/mL MTX before being plated on both selective (LB + MTX 32 mg/mL) and non-selective media. Clones isolated from all plates were re-streaked on non-selective media before being frozen down for characterization.

### Supplementary Tables

**Table S1:** List of bacterial strains used and constructed for this study.

| Strain | Genotype | Reference |
|---|---|---|
| DA4201/DA5438 | Wild type *E. coli* MG1655 (parent) | DA Strain Collection[1] |
| DA45134 | DA5438, ΔIS*150*::*yetiYFP* pSIM5-*tet* | DA Strain Collection[1] |
| DA52562 | DA45134, ΔIS*150*::*sacB-cat* | This study |
| DA52626 | DA52562, ΔIS*150*::*mTagBFP2* pSIM5-*tet* | This study |
| DA55343 | DA52562, ΔIS*150*::*sYFP2* pSIM5-*tet* | This study |
| DA55288 | DA52626, dupl[ΔIS*150*::*mTagBFP2*]**cat-sacB* pSIM5-*tet* | This study |
| DA55290 | DA55343, dupl[ΔIS*150*::*sYFP2*]**cat-sacB* pSIM5-*tet* | This study |
| DA56507 | DA4201, ΔIS*150*::*sYFP2* | This study |
| DA52848 | DA4201, ΔIS*150*::*mTagBFP2* | This study |
| MP18-03 | DA56507, C>T 58 bp upstream of *folA* | This study |
| MP18-04 | DA56507, W30R, C>T 58 bp upstream of *folA* | This study |
| MP18-05 | DA56507 pG06-VIM-1 | This study |
| MP18-06 | DA52848, C>T 58 bp upstream of *folA* | This study |
| MP18-07 | DA52848, W30R, C>T 58 bp upstream of *folA* | This study |
| MP18-08 | DA52848 pG06-VIM-1 | This study |
| MP18-09 | *E. coli* DH5α | Thermo Scientific |
| MP18-10 | MP18-09 pBAD30 | This study |
| MP18-11 | MP18-09 pBAD30_*dfrA1* | This study |
| MP18-12 | MP18-09 pBAD30_*dfrA12* | This study |
| MP06-01 | *E. coli* K56-2 ST73 | (*15*) |
| MP06-05 | MP06-01, W30R, C>T 58 bp upstream of *folA* | (*14*) |
| MP06-06 | *E. coli* K56-12 ST104 | (*15*) |
| MP06-11 | *E. coli* K56-16 ST127 | (*15*) |
| MP06-16 | *E. coli* K56-41 ST73 | (*15*) |
| MP06-21 | *E. coli* K56-44 ST12 | (*15*) |
| MP06-26 | *E. coli* K56-50 ST100 | (*15*) |
| MP06-31 | *E. coli* K56-68 ST95 | (*15*) |
| MP06-36 | *E. coli* K56-70 ST537 | (*15*) |
| MP06-41 | *E. coli* K56-75 ST69 | (*15*) |
| MP05-31 | MP06-41 p06-VIM-1 | (*16*) |
| MP06-46 | *E. coli* K56-78 ST1235 | (*15*) |
| MP18-13 | MP06-01, mutant isolated during MTX selective plating | This study |
| MP18-14 | MP06-01, mutant isolated during MTX selective plating | This study |
| MP18-15 | MP06-01, mutant isolated during MTX selective plating | This study |
| MP18-16 | MP06-01, mutant isolated during MTX selective plating | This study |
| MP18-17 | MP06-01, mutant isolated during MTX selective plating | This study |
| MP18-18 | MP06-01, mutant isolated during MTX selective plating | This study |
| MP18-19 | MP06-01, mutant isolated during MTX selective plating | This study |

| | | |
|---|---|---|
| MP18-20 | MP06-01, mutant isolated during MTX selective plating | This study |
| MP18-21 | MP06-01, mutant isolated during MTX selective plating | This study |
| MP18-22 | MP06-01, mutant isolated during MTX selective plating | This study |
| MP18-23 | MP06-01, mutant isolated during MTX selective plating | This study |
| MP18-24 | MP06-01, mutant isolated during MTX selective plating | This study |
| MP18-25 | MP06-01, mutant isolated during MTX selective plating | This study |
| MP18-26 | MP06-01, mutant isolated during MTX selective plating | This study |
| MP18-27 | MP06-01, mutant isolated during MTX selective plating | This study |
| MP18-28 | MP06-01, mutant isolated during MTX selective plating | This study |
| MP19-78 | MP06-01, mutant isolated during MTX sub-MIC evolution | This study |
| MP19-79 | MP06-01, mutant isolated during MTX sub-MIC evolution | This study |
| MP19-80 | MP06-01, mutant isolated during MTX sub-MIC evolution | This study |
| MP19-81 | MP06-01, mutant isolated during MTX sub-MIC evolution | This study |
| MP20-01 | MP06-01, mutant isolated during MTX sub-MIC evolution | This study |
| MP20-02 | MP06-01, mutant isolated during MTX sub-MIC evolution | This study |
| MP20-03 | MP06-01, mutant isolated during MTX sub-MIC evolution | This study |
| MP20-04 | MP06-01, mutant isolated during MTX sub-MIC evolution | This study |
| MP20-05 | MP06-01, mutant isolated during MTX sub-MIC evolution | This study |
| MP20-06 | MP06-01, mutant isolated during MTX sub-MIC evolution | This study |
| MP20-07 | MP06-01, mutant isolated during MTX sub-MIC evolution | This study |
| MP20-08 | MP06-01, mutant isolated during MTX sub-MIC evolution | This study |
| MP20-09 | MP06-01, mutant isolated during MTX sub-MIC evolution | This study |
| MP20-10 | MP06-01, mutant isolated during MTX sub-MIC evolution | This study |
| MP20-11 | MP06-01, mutant isolated during MTX sub-MIC evolution | This study |
| MP20-12 | MP06-01, mutant isolated during MTX sub-MIC evolution | This study |
| MP20-13 | MP06-01, mutant isolated during MTX sub-MIC evolution | This study |
| MP20-14 | MP06-01, mutant isolated during MTX sub-MIC evolution | This study |
| MP20-15 | MP06-01, mutant isolated during MTX sub-MIC evolution | This study |
| MP20-16 | MP06-01, mutant isolated during MTX sub-MIC evolution | This study |
| MP20-17 | MP06-01, mutant isolated during MTX sub-MIC evolution | This study |
| MP20-18 | MP06-01, mutant isolated during MTX sub-MIC evolution | This study |
| MP20-19 | MP06-01, mutant isolated during MTX sub-MIC evolution | This study |
| MP20-20 | MP06-01, mutant isolated during MTX sub-MIC evolution | This study |
| MP20-21 | MP06-01, mutant isolated during MTX sub-MIC evolution | This study |
| MP20-22 | MP06-01, mutant isolated during MTX sub-MIC evolution | This study |
| MP20-23 | MP06-01, mutant isolated during MTX sub-MIC evolution | This study |
| MP20-24 | MP06-01, mutant isolated during MTX sub-MIC evolution | This study |
| MP20-25 | MP06-01, mutant isolated during MTX sub-MIC evolution | This study |
| MP20-26 | MP06-01, mutant isolated during MTX sub-MIC evolution | This study |
| MP20-27 | MP06-01, mutant isolated during MTX sub-MIC evolution | This study |
| MP20-28 | MP06-01, mutant isolated during MTX sub-MIC evolution | This study |

| | | |
|---|---|---|
| MP20-29 | MP06-01, mutant isolated during MTX sub-MIC evolution | This study |
| MP20-30 | MP06-01, mutant isolated during MTX sub-MIC evolution | This study |
| MP20-31 | MP06-01, mutant isolated during MTX sub-MIC evolution | This study |
| MP20-32 | MP06-01, mutant isolated during MTX sub-MIC evolution | This study |
| MP20-33 | MP06-01, mutant isolated during MTX sub-MIC evolution | This study |
| MP20-34 | MP06-01, mutant isolated during MTX sub-MIC evolution | This study |
| MP20-35 | MP06-01, mutant isolated during MTX sub-MIC evolution | This study |
| MP20-36 | MP06-01, mutant isolated during MTX sub-MIC evolution | This study |
| MP20-37 | MP06-01, mutant isolated during MTX sub-MIC evolution | This study |
| MP20-38 | MP06-01, mutant isolated during MTX sub-MIC evolution | This study |
| MP20-39 | MP06-01, mutant isolated during MTX sub-MIC evolution | This study |
| MP20-40 | MP06-01, mutant isolated during MTX sub-MIC evolution | This study |
| MP20-41 | MP06-01, mutant isolated during MTX sub-MIC evolution | This study |
| MP20-42 | MP06-01, mutant isolated during MTX sub-MIC evolution | This study |

[1]DA strain collection refers to the bacterial strain collection in the Dan Andersson lab at Uppsala University.

**Table S2:** MTX and TMP susceptibility profiles of strains used and constructed for the study. Strains with known TMP resistance determinants have been highlighted as bold in the table. The final MIC concentration was set as the modal value from 3-5 replicates (Table S15).

| **Strain** | **MTX MIC [mg/mL]** | **TMP MIC [µg/mL]** |
|---|---|---|
| DA4201/DA5438 | 16 | 0.125 |
| DA56507 | 16 | 0.125 |
| DA52848 | 8 | 0.25 |
| **MP18-03** | **>32** | **4** |
| **MP18-04** | **>32** | **32** |
| **MP18-05** | **>32** | **>64** |
| **MP18-06** | **>32** | **8** |
| **MP18-07** | **>32** | **32** |
| **MP18-08** | **>32** | **>64** |
| MP18-09 | <2 | 0.0625 |
| MP18-10 | <2 | 0.0625 |
| **MP18-11**[3] | **<2** | **0.0625** |
| **MP18-12** | **>32** | **>64** |
| MP06-01[1, 2] | 4 | 0.225 |
| **MP06-05** | **>32** | **≥28**[2] |
| MP06-06 | 32 | 0.563[2] |
| MP06-11 | 4 | 0.250[2] |
| MP06-16 | 16 | 0.250[2] |
| MP06-21 | 4 | 0.375[2] |
| MP06-26 | 8 | 0.172[2] |
| MP06-31 | 32 | 0.208[2] |
| MP06-36 | 4 | 0.250[2] |
| MP06-41 | 4 | 0.167[2] |
| **MP05-31** | **>32** | **>64** |
| MP06-46 | 32 | 0.500[2] |

[1]MP06-01 used as internal standard on all MXT-plates, modal MIC based on 50 replicates.

[2]TMP MICs determined in (*14*).

[3]MP18-11 has been marked in bold as the strain expresses the known $\text{TMP}^R$ determinant *dfrA*1. The strain did however not confer resistance to either MTX or TMP.

**Table S3:** Calculated selection coefficients of each replicate during head-to-head competitions between the TMP$^{R}$ *folA* mutant (MP18-04 and MP18-07) and isogenic TMP$^{S}$ (DA56507 and DA52848) strain.

| **MTX concentration [µg/mL]** | **0** | **200** | **400** | **600** | **800** | **1,000** |
|---|---|---|---|---|---|---|
| Selection coefficients *bfp*R:*yfp*S (MP18-07:DA56507) | -0.0135075 | -0.0044673 | -0.0020299 | 0.03931029 | 0.03190433 | 0.07611339 |
| | -0.0255208 | -0.0067445 | 0.00602582 | 0.03781498 | 0.04498202 | 0.08812079 |
| | -0.0388344 | -0.0005859 | 0.00986104 | 0.0355561 | 0.07659162 | 0.09771619 |
| | -0.0320059 | -0.0023467 | 0.00807802 | 0.04331207 | 0.05821313 | 0.1132978 |
| | -0.0364165 | -0.0057047 | 0.00550519 | 0.0347153 | 0.04307795 | 0.12888063 |
| | -0.0255894 | -0.0079951 | 0.01038676 | 0.01334624 | 0.04880345 | 0.09785017 |
| Selection coefficients *yfp*R:*bfp*S (MP18-04:DA52848) | -0.0372175 | -0.0005588 | 0.02263352 | 0.08257795 | 0.19266 | 0.1671059 |
| | -0.0210616 | 0.00438223 | 0.04124424 | 0.12591823 | 0.15959803 | 0.31567517 |
| | -0.0316734 | 0.0055524 | 0.03384889 | 0.12513247 | 0.1767099 | 0.26984198 |
| | -0.0315623 | 0.00256964 | 0.04051869 | 0.08922083 | 0.20218262 | 0.24558123 |
| | -0.0317976 | 0.00157104 | 0.02518501 | 0.09396005 | 0.18303477 | 0.19802551 |
| | -0.0356574 | 0.00039645 | 0.04996337 | 0.09123521 | 0.1754452 | 0.25601346 |
| Mean | -0.0300703 | -0.0011609 | 0.02093506 | 0.06767498 | 0.11610025 | 0.17118518 |
| St. dv. | 0.00711319 | 0.00419485 | 0.01636382 | 0.03649801 | 0.06693214 | 0.07954708 |
| N | 12 | 12 | 12 | 12 | 12 | 12 |

**Table S4:** Calculated selection coefficients of each replicate during head-to-head competitions between the TMP$^{R}$ *E. coli* MG1655 harboring the pG06-VIM-1 plasmid (MP18-05 and MP18-08) and isogenic TMP$^{S}$ strain (DA56507 and DA52848).

| **MTX concentration [µg/mL]** | **0** | **25** | **50** | **75** | **100** | **200** |
|---|---|---|---|---|---|---|
| Selection coefficients *bfp*R:*yfp*S (MP18-08:DA56507) | 0.0013778 | -0.0010507 | 0.0055619 | 0.01118738 | 0.01059141 | 0.02713763 |
| | 0.00063436 | 0.00289697 | 0.00504481 | 0.01138153 | 0.01172904 | 0.03061104 |
| | 0.00058383 | 0.00169671 | 0.00429196 | 0.00686043 | 0.01459478 | 0.03452533 |
| | 0.0014676 | 0.0016126 | 0.00519585 | 0.01026467 | 0.01410065 | 0.02326364 |
| | -0.0007107 | -0.0004971 | 0.00718521 | 0.00893631 | 0.00907501 | 0.03498757 |
| | 0.00111876 | 0.0019667 | 0.00499255 | 0.01192992 | 0.01541848 | 0.03323181 |
| Selection coefficients *yfp*R:*bfp*S (MP18-05:DA52848) | 0.00370853 | 0.01542282 | 0.00627514 | 0.00868095 | 0.0212524 | 0.04832319 |
| | 0.00665488 | 0.00014057 | 0.00940239 | 0.02035904 | 0.02432032 | 0.06098507 |
| | 0.00594351 | 0.00041042 | 0.01254878 | 0.01541574 | 0.02045735 | 0.06160274 |
| | 0.00515497 | 0.0072364 | 0.01024738 | 0.01097359 | 0.02139386 | 0.07722027 |
| | 0.00331283 | 0.00814149 | 0.01039754 | 0.01232684 | 0.01975182 | 0.04729734 |
| | 0.00567272 | 0.0017711 | 0.00961744 | 0.01822696 | 0.01961757 | 0.05169943 |
| Mean | 0.00290992 | 0.00331234 | 0.00756341 | 0.01221195 | 0.01685856 | 0.04424042 |
| St. dv. | 0.00238108 | 0.00453936 | 0.00262615 | 0.00378389 | 0.00470341 | 0.01573381 |
| N | 12 | 12 | 12 | 12 | 12 | 12 |

**Table S5:** Isogenic TMP$^{R}$ and TMP$^{S}$ *E. coli* MG1655 strains were competed against each other in six biological replicates, at different starting ratios and changes in *bfp*R:*yfp*S ratios over 30 generations were measured. Replicates where the starting ratio was not as intended and where random mutations have occurred during the competition experiment (reflected as a nonlinear slope) have been excluded from the calculations (*18*).

| TMP$^{R}$ | MTX concentration | Replicate | Starting ratio | Ratio after 10 generations | Ratio after 20 generations | Ratio after 30 generations |
|---|---|---|---|---|---|---|
| *folA* | 400 µg/mL | 1 | 1 | 0.990 | 1.127 | 1.317 |
| | | 2 | 1 | 0.927 | 1.099 | 1.096 |
| | | 3 | 1 | 0.995 | 1.215 | 1.292 |
| | | 4 | 1 | 0.949 | 1.115 | 1.097 |
| | | 5 | 1 | 0.984 | 1.072 | 1.091 |
| | | 1 | 0.1 | 0.535 | 0.711 | 0.738 |
| | | 2 | 0.1 | 0.583 | 0.741 | 0.774 |
| | | 3 | 0.1 | 0.615 | 0.758 | 0.839 |
| | | 4 | 0.1 | 0.607 | 0.734 | 0.760 |
| | | 5 | 0.1 | 0.606 | 0.745 | 0.789 |
| | | 1 | 0.01 | 0.127 | 0.319 | 0.530 |
| | | 2 | 0.01 | 0.154 | 0.347 | 0.456 |
| | | 3 | 0.01 | 0.153 | 0.390 | 0.453 |
| | | 4 | 0.01 | 0.120 | 0.333 | 0.375 |
| | | 5 | 0.01 | 0.171 | 0.316 | 0.446 |
| | | 1 | 0.001 | 0.029 | 0.105 | 0.174 |
| | | 2 | 0.001 | 0.015 | 0.037 | 0.093 |
| | | 3 | 0.001 | 0.023 | 0.068 | 0.117 |
| | | 4 | 0.001 | 0.023 | 0.053 | 0.079 |
| | | 5 | 0.001 | 0.027 | 0.056 | 0.102 |
| | | 1 | 0.0001 | 0.002 | 0.004 | 0.006 |
| | | 2 | 0.0001 | 0.002 | 0.005 | 0.007 |
| | | 3 | 0.0001 | 0.003 | 0.008 | 0.015 |
| | | 5 | 0.0001 | 0.003 | 0.006 | 0.007 |
| pG06-VIM-1 | 75 µg/mL | 1 | 1 | 0.927 | 1.017 | 1.257 |
| | | 2 | 1 | 1.054 | 1.152 | 1.296 |
| | | 3 | 1 | 0.977 | 1.100 | 1.264 |
| | | 4 | 1 | 1.002 | 1.095 | 1.297 |
| | | 6 | 1 | 0.880 | 0.979 | 1.087 |
| | | 1 | 0.1 | 0.490 | 0.670 | 0.772 |
| | | 2 | 0.1 | 0.536 | 0.702 | 0.801 |
| | | 3 | 0.1 | 0.490 | 0.678 | 0.771 |
| | | 4 | 0.1 | 0.644 | 0.744 | 0.816 |
| | | 6 | 0.1 | 0.627 | 0.774 | 0.813 |
| | | 1 | 0.01 | 0.091 | 0.161 | 0.301 |
| | | 2 | 0.01 | 0.115 | 0.200 | 0.304 |
| | | 3 | 0.01 | 0.124 | 0.230 | 0.258 |
| | | 4 | 0.01 | 0.148 | 0.242 | 0.346 |
| | | 6 | 0.01 | 0.132 | 0.202 | 0.325 |
| | | 1 | 0.001 | 0.009 | 0.016 | 0.035 |
| | | 2 | 0.001 | 0.018 | 0.030 | 0.057 |
| | | 3 | 0.001 | 0.007 | 0.014 | 0.019 |
| | | 4 | 0.001 | 0.012 | 0.021 | 0.033 |

| | | | | |
|---|---|---|---|---|
| 6 | 0.001 | 0.016 | 0.028 | 0.041 |
| 1 | 0.0001 | 0.001 | 0.001 | 0.004 |
| 2 | 0.0001 | 0.002 | 0.003 | 0.007 |
| 3 | 0.0001 | 0.002 | 0.004 | 0.004 |
| 5 | 0.0001 | 0.002 | 0.004 | 0.005 |
| 6 | 0.0001 | 0.001 | 0.002 | 0.005 |

**Table S6:** List of clones isolated during selective plating on high concentrations of MTX, including MTX and TMP susceptibility profiles as well as results from Sanger sequencing of the *folA* gene, its promotor region and of the *marR* gene. The final MIC concentration was set as the modal value from 3-4 replicates (Table S15). See Protocols and Figure S2 for more detailed description on selection procedures.

| Strain | [MTX] in O.N. culture (mg/mL) | [MTX] in plates (mg/mL) | MTX MIC (mg/mL) | TMP MIC (µg/mL) | Mutations in *folA*/$P_{folA}$ | Mutations in *marR* |
|---|---|---|---|---|---|---|
| MP18-13 | 0 | 4 | 8 | <0.03125 | None | None |
| MP18-14 | 0 | 4 | 32 | 0.125 | None | None |
| MP18-15 | 0 | 8 | 4 | 0.125 | None | None |
| MP18-16 | 0 | 8 | >32 | 0.125 | None | None |
| MP18-17 | 0 | 16 | >32 | >64 | G>A 32 bp upstream | None |
| MP18-18 | 0 | 16 | >32 | 2 | G>A 32 bp upstream | None |
| MP18-19 | 0 | 16 | >32 | 4 | G>A 32 bp upstream | None |
| MP18-20 | 6.25 | 32 | >32 | 4 | C>T 58 bp upstream | A41E |
| MP18-21 | 6.25 | 32 | >32 | 4 | C>T 58 bp upstream | A41E |
| MP18-22 | 6.25 | 32 | >32 | 4 | C>T 58 bp upstream | A41E |
| MP18-23 | 6.25 | 32 | >32 | 8 | C>T 58 bp upstream | A41E |
| MP18-24 | 6.25 | 32 | >32 | 4 | C>T 58 bp upstream | A41E |
| MP18-25 | 6.25 | 32 | >32 | 4 | C>T 58 bp upstream | A41E |
| MP18-26 | 6.25 | 32 | >32 | 4 | C>T 58 bp upstream | A41E |
| MP18-27 | 6.25 | 32 | >32 | 4 | C>T 58 bp upstream | A41E |
| MP18-28 | 6.25 | 0 | >32 | 4 | C>T 58 bp upstream | A41E |

**Table S7:** Putative mutations identified during WGS analysis of MP18-13 when compared to the *E. coli* K56-2 WT strain used for selective plating. No mutations were found in genes known to play a role in antibiotic resistance.

| Mutation type | Original base | Alternative base | Evidence | Strand | Effect | Gene | Product |
|---|---|---|---|---|---|---|---|
| SNP | G | T | T:605 G:0 | + | Asp577Tyr | | gamma-glutamyltransferase |
| SNP | T | C | C:523 T:0 | + | Synonymous variant Phe411Phe | *proY* | proline-specific permease ProY |
| Deletion | CA | C | C:343 CA:0 | | | | |
| Insertion | A | AT | AT:405 A:0 | | | | |
| SNP | T | C | C:417 T:0 | | | | |
| SNP | A | C | C:634 A:0 | - | Cys594Gly | | beta-galactosidase |
| SNP | G | T | T:610 G:0 | + | Asp241Tyr | | 3-phenylpropionate MFS transporter |
| SNP | G | C | C:635 G:0 | - | Arg93Gly | *arnC* | undecaprenyl-phosphate 4-deoxy-4-formamido-L-arabinose transferase |
| SNP | G | C | C:629 G:0 | - | Leu91Val | *arnC* | undecaprenyl-phosphate 4-deoxy-4-formamido-L-arabinose transferase |
| SNP | C | T | T:635 C:0 | - | Synonymous variant Gly87Gly | *arnC* | undecaprenyl-phosphate 4-deoxy-4-formamido-L-arabinose transferase |
| SNP | G | C | C:713 G:0 | + | Synonymous variant Gly289Gly | *yfaL* | AIDA-I family autotransporter adhesin YfaL/EhaC |
| SNP | T | A | A:576 T:0 | - | Stop lost Ter116Tyrext*? | *yejF* | microcin C ABC transporter ATP-binding protein YejF |
| SNP | T | G | G:728 T:2 | + | Synonymous variant Pro229Pro | | DEAD/DEAH box helicase family protein |
| SNP | A | G | G:712 A:52 | + | Met2Val | | IS66 family transposase |
| Insertion | G | GA | GA:631 G:0 | + | Frameshift variant & stop lost Ter366fs | | kfiB protein |
| SNP | G | C | C:552 G:0 | + | Synonymous variant Gly403Gly | | purine permease |
| SNP | G | A | A:624 G:0 | + | Val115Ile | *nimT* | 2-nitroimidazole transporter |
| SNP | C | A | A:757 C:0 | + | Pro3Thr | | biofilm-dependent modulation protein |
| SNP | C | A | A:609 C:0 | + | Ser557Tyr | *maeA* | oxaloacetate-decarboxylating malate dehydrogenase |
| Complex | CCCC | ACCT | ACCT:617 CCCC:0 | - | Synonymous variant | | ABC transporter permease |
| SNP | C | T | T:678 C:0 | - | Val136Met | *tehB* | tellurite resistance methyltransferase TehB |
| SNP | G | A | A:287 G:23 | - | Synonymous variant Gly315Gly | | EntS/YbdA MFS transporter |
| SNP | T | A | A:636 T:2 | + | Phe106Leu | *pdeR* | cyclic di-GMP phosphodiesterase |
| SNP | A | C | C:664 A:0 | + | Asn306His | *pdeR* | cyclic di-GMP phosphodiesterase |
| SNP | G | T | T:620 G:0 | - | Asp56Glu | *tonB* | TonB system transport protein TonB |
| SNP | T | C | C:603 T:0 | - | Synonymous variant Leu366Leu | | autotransporter outer membrane beta-barrel domain-containing protein |
| Deletion | TC | T | T:630 TC:0 | - | Frameshift variant Asp86fs | *rfaP* | lipopolysaccharide core heptose(I) kinase RfaP |
| SNP | G | A | A:629 G:0 | - | Synonymous variant Asn433Asn | *ilvB* | acetolactate synthase large subunit |
| SNP | C | A | A:683 C:0 | - | Gly2778Val | *clbB* | colibactin hybrid non-ribosomal peptide synthetase/type I polyketide synthase ClbB |

| | | | | | | | |
|---|---|---|---|---|---|---|---|
| SNP | G | T | T:622 G:0 | + | Synonymous variant Gly538Gly | *ftsI* | peptidoglycan glycosyltransferase FtsI |

**Table S8:** Putative mutations identified during WGS analysis of MP18-17 when compared to the *E. coli* K56-2 WT strain used for selective plating. The strain was shown to have a G > A mutation 32 base pairs upstream of the *folA* gene, in its promotor, whereas no other mutations were found in genes known to play a role in antibiotic resistance.

| Mutation type | Original base | Alternative base | Evidence | Strand | Effect | Gene | Product |
|---|---|---|---|---|---|---|---|
| SNP | G | T | T:601 G:0 | + | Asp577Tyr | | gamma-glutamyltransferase |
| SNP | T | C | C:490 T:0 | + | Synonymous variant Phe411Phe | *proY* | proline-specific permease ProY |
| Deletion | CA | C | C:360 CA:0 | | | | |
| Insertion | A | AT | AT:450 A:0 | | | | |
| SNP | T | C | C:384 T:0 | | | | |
| SNP | A | C | C:602 A:0 | - | Cys594Gly | | beta-galactosidase |
| SNP | G | T | T:513 G:0 | + | Asp241Tyr | | 3-phenylpropionate MFS transporter |
| SNP | G | C | C:635 G:0 | - | Arg93Gly | *arnC* | undecaprenyl-phosphate 4-deoxy-4-formamido-L-arabinose transferase |
| SNP | G | C | C:632 G:0 | - | Leu91Val | *arnC* | undecaprenyl-phosphate 4-deoxy-4-formamido-L-arabinose transferase |
| SNP | C | T | T:647 C:0 | - | Synonymous variant Gly87Gly | *arnC* | undecaprenyl-phosphate 4-deoxy-4-formamido-L-arabinose transferase |
| SNP | G | C | C:690 G:0 | + | Synonymous variant Gly289Gly | *yfaL* | AIDA-I family autotransporter adhesin YfaL/EhaC |
| SNP | T | A | A:547 T:0 | - | Stop lost Ter116Tyrext*? | *yejF* | microcin C ABC transporter ATP-binding protein YejF |
| SNP | A | C | C:735 A:0 | + | Asn75His | | glycosyltransferase family 2 protein |
| SNP | T | G | G:604 T:0 | + | Synonymous variant Pro229Pro | | DEAD/DEAH box helicase family protein |
| SNP | A | G | G:663 A:63 | + | Met2Val | | IS66 family transposase |
| Insertion | G | GA | GA:558 G:1 | + | Frameshift variant & stop lost Ter366fs | | kfiB protein |
| SNP | G | C | C:555 G:0 | + | Synonymous variant Gly403Gly | | purine permease |
| SNP | G | A | A:572 G:0 | + | Val115Ile | *nimT* | 2-nitroimidazole transporter |
| SNP | C | A | A:800 C:0 | + | Pro3Thr | | biofilm-dependent modulation protein |
| SNP | C | A | A:606 C:0 | + | Ser557Tyr | *maeA* | oxaloacetate-decarboxylating malate dehydrogenase |
| Complex | CCCC | ACCT | ACCT:525 CCCC:0 | - | Synonymous variant | | ABC transporter permease |
| SNP | C | T | T:651 C:0 | - | Val136Met | *tehB* | tellurite resistance methyltransferase TehB |
| SNP | G | A | A:295 G:18 | - | Synonymous variant Gly315Gly | | EntS/YbdA MFS transporter |
| SNP | T | A | A:622 T:2 | + | Phe106Leu | *pdeR* | cyclic di-GMP phosphodiesterase |
| SNP | A | C | C:564 A:0 | + | Asn306His | *pdeR* | cyclic di-GMP phosphodiesterase |
| SNP | G | T | T:496 G:0 | - | Asp56Glu | *tonB* | TonB system transport protein TonB |
| SNP | T | C | C:554 T:0 | - | Synonymous variant Leu366Leu | | autotransporter outer membrane beta-barrel domain-containing protein |

| | | | | | | | |
|---|---|---|---|---|---|---|---|
| SNP | G | A | A:532 G:0 | - | Synonymous variant Asn433Asn | *ilvB* | acetolactate synthase large subunit |
| SNP | C | A | A:585 C:0 | - | Gly2778Val | *clbB* | colibactin hybrid non-ribosomal peptide synthetase/type I polyketide synthase ClbB |
| SNP | G | A | A:643 G:0 | | | Promotor region of *folA* | dihydrofolate reductase *folA* |
| SNP | G | T | T:638 G:0 | + | Synonymous variant Gly538Gly | *ftsI* | peptidoglycan glycosyltransferase FtsI |

**Table S9:** Putative mutations identified during WGS analysis of MP18-20 when compared to the *E. coli* K56-2 WT strain used for selective plating. The strain was shown to have a C>A mutation in the *marR* gene (A41G) and a C > T mutation 58 base pairs upstream of the *folA* gene, in its promotor. No other mutations were found in genes known to play a role in antibiotic resistance.

| Mutation type | Original base | Alternative base | Evidence | Strand | Effect | Gene | Product |
|---|---|---|---|---|---|---|---|
| SNP | G | T | T:643 G:0 | + | Asp577Tyr | | gamma-glutamyltransferase |
| SNP | T | C | C:613 T:1 | + | Synonymous variant Phe411Phe | *proY* | proline-specific permease ProY |
| Deletion | CA | C | C:342 CA:0 | | | | |
| Insertion | A | AT | AT:398 A:1 | | | | |
| SNP | T | C | C:446 T:0 | | | | |
| SNP | A | C | C:636 A:0 | - | Cys594Gly | | beta-galactosidase |
| SNP | G | T | T:579 G:0 | + | Asp241Tyr | | 3-phenylpropionate MFS transporter |
| SNP | G | C | C:638 G:0 | - | Arg93Gly | *arnC* | undecaprenyl-phosphate 4-deoxy-4-formamido-L-arabinose transferase |
| SNP | G | C | C:640 G:0 | - | Leu91Val | *arnC* | undecaprenyl-phosphate 4-deoxy-4-formamido-L-arabinose transferase |
| SNP | C | T | T:645 C:0 | - | Synonymous variant Gly87Gly | *arnC* | undecaprenyl-phosphate 4-deoxy-4-formamido-L-arabinose transferase |
| SNP | G | C | C:740 G:0 | + | Synonymous variant Gly289Gly | *yfaL* | AIDA-I family autotransporter adhesin YfaL/EhaC |
| SNP | T | A | A:606 T:0 | - | Stop lost Ter116Tyrext*? | *yejF* | microcin C ABC transporter ATP-binding protein YejF |
| SNP | T | G | G:652 T:0 | + | Synonymous variant Pro229Pro | | DEAD/DEAH box helicase family protein |
| SNP | A | C | C:680 A:0 | - | Phe148Cys | | phosphoethanolamine transferase |
| SNP | A | G | G:726 A:66 | + | Met2Val | | IS66 family transposase |
| Insertion | G | GA | GA:686 G:0 | + | Frameshift variant & stop lost Ter366fs | | kfiB protein |
| SNP | G | C | C:598 G:0 | + | Synonymous variant Gly403Gly | | purine permease |
| SNP | C | A | A:736 C:0 | + | Ala41Glu | *marR* | multiple antibiotic resistance transcriptional regulator MarR |
| SNP | G | A | A:580 G:0 | + | Val115Ile | *nimT* | 2-nitroimidazole transporter |
| SNP | C | A | A:895 C:0 | + | Pro3Thr | | biofilm-dependent modulation protein |

| | | | | | | | |
|---|---|---|---|---|---|---|---|
| SNP | C | A | A:710 C:0 | + | Ser557Tyr | *maeA* | oxaloacetate-decarboxylating malate dehydrogenase |
| Complex | CCCC | ACCT | ACCT:585 CCCC:0 | - | Synonymous variant | | ABC transporter permease |
| SNP | C | T | T:787 C:0 | - | Val136Met | *tehB* | tellurite resistance methyltransferase TehB |
| SNP | G | A | A:331 G:15 | - | Synonymous variant Gly315Gly | | EntS/YbdA MFS transporter |
| SNP | T | A | A:674 T:0 | + | Phe106Leu | *pdeR* | cyclic di-GMP phosphodiesterase |
| SNP | A | C | C:630 A:0 | + | Asn306His | *pdeR* | cyclic di-GMP phosphodiesterase |
| SNP | G | T | T:595 G:0 | - | Asp56Glu | *tonB* | TonB system transport protein TonB |
| SNP | T | C | C:564 T:0 | - | Synonymous variant Leu366Leu | | autotransporter outer membrane beta-barrel domain-containing protein |
| SNP | C | A | A:666 C:0 | - | Gly2778Val | *clbB* | colibactin hybrid non-ribosomal peptide synthetase/type I polyketide synthase ClbB |
| SNP | C | T | T:697 C:0 | | | Promotor region of *folA* | dihydrofolate reductase *folA* |
| SNP | G | T | T:626 G:0 | + | Synonymous variant Gly538Gly | *ftsI* | peptidoglycan glycosyltransferase FtsI |

**Table S10:** Putative mutations identified during WGS analysis of MP18-26 when compared to the *E. coli* K56-2 WT strain used for selective plating. The strain was shown to have a C>A mutation in the *marR* gene (A41G) and a C > T mutation 58 base pairs upstream of the *folA* gene, in its promotor. No other mutations were found in genes known to play a role in antibiotic resistance.

| Mutation type | Original base | Alternative base | Evidence | Strand | Effect | Gene | Product |
|---|---|---|---|---|---|---|---|
| SNP | G | T | T:620 G:0 | + | Asp577Tyr | | gamma-glutamyltransferase |
| SNP | T | C | C:527 T:0 | + | Synonymous variant Phe411Phe | *proY* | proline-specific permease ProY |
| Deletion | CA | C | C:318 CA:0 | | | | |
| Insertion | A | AT | AT:360 A:0 | | | | |
| SNP | T | C | C:337 T:0 | | | | |
| SNP | A | C | C:565 A:0 | - | Cys594Gly | | beta-galactosidase |
| SNP | G | T | T:475 G:0 | + | Asp241Tyr | | 3-phenylpropionate MFS transporter |
| SNP | G | C | C:580 G:0 | - | Arg93Gly | *arnC* | undecaprenyl-phosphate 4-deoxy-4-formamido-L-arabinose transferase |
| SNP | G | C | C:573 G:0 | - | Leu91Val | *arnC* | undecaprenyl-phosphate 4-deoxy-4-formamido-L-arabinose transferase |
| SNP | C | T | T:577 C:0 | - | Synonymous variant Gly87Gly | *arnC* | undecaprenyl-phosphate 4-deoxy-4-formamido-L-arabinose transferase |
| SNP | G | C | C:628 G:0 | + | Synonymous variant Gly289Gly | *yfaL* | AIDA-I family autotransporter adhesin YfaL/EhaC |
| SNP | T | A | A:526 T:1 | - | Stop lost Ter116Tyrext*? | *yejF* | microcin C ABC transporter ATP-binding protein YejF |

| | | | | | | | |
|---|---|---|---|---|---|---|---|
| SNP | T | G | G:572 T:0 | + | Synonymous variant Pro229Pro | | DEAD/DEAH box helicase family protein |
| SNP | A | C | C:607 A:0 | - | Phe148Cys | | phosphoethanolamine transferase |
| SNP | A | G | G:581 A:45 | + | Met2Val | | IS66 family transposase |
| Insertion | G | GA | GA:542 G:1 | + | Frameshift variant & stop lost Ter366fs | | kfiB protein |
| SNP | G | C | C:525 G:0 | + | Synonymous variant Gly403Gly | | purine permease |
| SNP | C | A | A:649 C:0 | + | Ala41Glu | *marR* | multiple antibiotic resistance transcriptional regulator MarR |
| SNP | G | A | A:570 G:0 | + | Val115Ile | *nimT* | 2-nitroimidazole transporter |
| SNP | C | A | A:715 C:0 | + | Pro3Thr | | biofilm-dependent modulation protein |
| SNP | C | A | A:561 C:0 | + | Ser557Tyr | *maeA* | oxaloacetate-decarboxylating malate dehydrogenase |
| Complex | CCCC | ACCT | ACCT:427 CCCC:0 | - | Synonymous variant | | ABC transporter permease |
| SNP | C | T | T:655 C:0 | - | Val136Met | *tehB* | tellurite resistance methyltransferase TehB |
| SNP | T | A | A:576 T:1 | + | Phe106Leu | *pdeR* | cyclic di-GMP phosphodiesterase |
| SNP | A | C | C:588 A:0 | + | Asn306His | *pdeR* | cyclic di-GMP phosphodiesterase |
| SNP | G | T | T:597 G:0 | - | Asp56Glu | *tonB* | TonB system transport protein TonB |
| SNP | T | C | C:511 T:0 | - | Synonymous variant Leu366Leu | | autotransporter outer membrane beta-barrel domain-containing protein |
| SNP | C | A | A:548 C:0 | - | Gly2778Val | *clbB* | colibactin hybrid non-ribosomal peptide synthetase/type I polyketide synthase ClbB |
| SNP | C | T | T:527 C:0 | | | Promotor region of *folA* | dihydrofolate reductase *folA* |
| SNP | G | T | T:557 G:0 | + | Synonymous variant Gly538Gly | *ftsI* | peptidoglycan glycosyltransferase FtsI |

**Table S11:** Putative mutations identified during WGS analysis of MP18-28 when compared to the *E. coli* K56-2 WT strain used for selective plating. The strain was shown to have a C>A mutation in the *marR* gene (A41G) and a C > T mutation 58 base pairs upstream of the *folA* gene, in its promotor. No other mutations were found in genes known to play a role in antibiotic resistance.

| **Mutation type** | **Original base** | **Alternative base** | **Evidence** | **Strand** | **Effect** | **Gene** | **Product** |
|---|---|---|---|---|---|---|---|
| SNP | G | T | T:370 G:0 | + | Asp577Tyr | | gamma-glutamyltransferase |
| SNP | T | C | C:371 T:0 | + | Synonymous variant Phe411Phe | *proY* | proline-specific permease ProY |
| Deletion | CA | C | C:208 CA:0 | | | | |
| ins | A | AT | AT:282 A:0 | | | | |
| SNP | T | C | C:238 T:0 | | | | |
| SNP | A | C | C:427 A:0 | - | Cys594Gly | | beta-galactosidase |
| SNP | G | T | T:320 G:0 | + | Asp241Tyr | | 3-phenylpropionate MFS transporter |
| SNP | G | C | C:370 G:0 | - | Arg93Gly | *arnC* | undecaprenyl-phosphate 4-deoxy-4- |

| | | | | | | | |
|---|---|---|---|---|---|---|---|
| | | | | | | | formamido-L- arabinose transferase |
| SNP | G | C | C:366 G:0 | - | Leu91Val | *arnC* | undecaprenyl- phosphate 4-deoxy-4- formamido-L- arabinose transferase |
| SNP | C | T | T:358 C:0 | - | Synonymous variant Gly87Gly | *arnC* | undecaprenyl- phosphate 4-deoxy-4- formamido-L- arabinose transferase |
| SNP | G | C | C:431 G:0 | + | Synonymous variant Gly289Gly | *yfaL* | AIDA-I family autotransporter adhesin YfaL/EhaC |
| SNP | T | A | A:332 T:0 | - | Stop lost Ter116Tyrext*? | *yejF* | microcin C ABC transporter ATP- binding protein YejF |
| SNP | T | G | G:399 T:0 | + | Synonymous variant Pro229Pro | | DEAD/DEAH box helicase family protein |
| SNP | A | C | C:423 A:0 | - | Phe148Cys | | phosphoethanolamine transferase |
| ins | G | GA | GA:366 G:0 | + | Frameshift variant & stop lost Ter366fs | | kfiB protein |
| SNP | G | C | C:402 G:0 | + | Synonymous variant Gly403Gly | | purine permease |
| SNP | C | A | A:419 C:0 | + | Ala41Glu | *marR* | multiple antibiotic resistance transcriptional regulator MarR |
| SNP | G | A | A:389 G:0 | + | Val115Ile | *nimT* | 2-nitroimidazole transporter |
| SNP | C | A | A:445 C:0 | + | Pro3Thr | | biofilm-dependent modulation protein |
| SNP | C | A | A:402 C:0 | + | Ser557Tyr | *maeA* | oxaloacetate- decarboxylating malate dehydrogenase |
| Complex | CCCC | ACCT | ACCT:381 CCCC:0 | - | Synonymous variant | | ABC transporter permease |
| SNP | C | T | T:382 C:0 | - | Val136Met | *tehB* | tellurite resistance methyltransferase TehB |
| SNP | G | A | A:210 G:10 | - | Gly315Gly | | EntS/YbdA MFS transporter |
| SNP | T | A | A:396 T:0 | + | Phe106Leu | *pdeR* | cyclic di-GMP phosphodiesterase |
| SNP | A | C | C:402 A:0 | + | Asn306His | *pdeR* | cyclic di-GMP phosphodiesterase |
| SNP | G | T | T:368 G:0 | - | Asp56Glu | *tonB* | TonB system transport protein TonB |
| SNP | T | C | C:377 T:0 | - | Synonymous variant Leu366Leu | | autotransporter outer membrane beta-barrel domain-containing protein |
| SNP | C | A | A:409 C:0 | - | Gly2778Val | *clbB* | colibactin hybrid non- ribosomal peptide synthetase/type I polyketide synthase ClbB |
| SNP | C | T | T:421 C:0 | | | Promotor region of *folA* | dihydrofolate reductase *folA* |
| SNP | G | T | T:422 G:0 | + | Synonymous variant Gly538Gly | *ftsI* | peptidoglycan glycosyltransferase FtsI |

**Table S12:** List of clones isolated on plates containing TMP 4 μg/mL during laboratory evolution of K56-2 (MP06-01) at 400 μg/mL MTX, including TMP MICs. The final MIC concentration was set as the modal value from 2-4 replicates. All isolates were found to have a single mutation (C>T) 58 bp upstream of the *folA* gene, in the *folA* promotor. Each biological replicate is represented by B and the replicate number.

| Strain | Isolated from | | TMP MIC [μg/mL] | | | | |
|---|---|---|---|---|---|---|---|
| | Generation | Lineage | B1 | B2 | B3 | B4 | Modal |
| MP19-78 | 50 | 1 | 8 | 12 | 12 | | 12 |
| MP19-79 | 50 | 1 | 12 | 8 | 12 | | 12 |
| MP19-80 | 50 | 1 | 8 | 16 | 16 | | 16 |
| MP19-81 | 50 | 1 | 8 | 12 | 12 | | 12 |
| MP20-01 | 50 | 1 | 12 | 12 | | | 12 |
| MP20-02 | 50 | 6 | 8 | 12 | 12 | | 12 |
| MP20-03 | 50 | 6 | 12 | 12 | | | 12 |
| MP20-04 | 50 | 9 | 12 | 12 | | | 12 |
| MP20-05 | 50 | 9 | 8 | 12 | 8 | | 8 |
| MP20-06 | 50 | 9 | 6 | 12 | 12 | | 12 |
| MP20-07 | 50 | 9 | 8 | 8 | | | 8 |
| MP20-08 | 50 | 9 | 6 | 12 | 12 | | 12 |
| MP20-09 | 100 | 1 | 12 | 12 | | | 12 |
| MP20-10 | 100 | 1 | 8 | 12 | 12 | | 12 |
| MP20-11 | 100 | 1 | 8 | 8 | | | 8 |
| MP20-12 | 100 | 1 | 12 | 4 | 12 | | 12 |
| MP20-13 | 100 | 1 | 12 | 12 | | | 12 |
| MP20-14 | 100 | 9 | 8 | 12 | 12 | | 12 |
| MP20-15 | 100 | 9 | 12 | 12 | | | 12 |
| MP20-16 | 100 | 9 | 12 | 12 | | | 12 |
| MP20-17 | 100 | 9 | 12 | 8 | 12 | | 12 |
| MP20-18 | 100 | 9 | 8 | 8 | | | 8 |
| MP20-19 | 150 | 1 | 8 | 8 | | | 8 |
| MP20-20 | 150 | 1 | 8 | 12 | 16 | 12 | 12 |
| MP20-21 | 150 | 1 | 8 | 8 | | | 8 |
| MP20-22 | 150 | 1 | 8 | 8 | | | 8 |
| MP20-23 | 150 | 1 | 8 | 8 | | | 8 |
| MP20-24 | 150 | 9 | 8 | 8 | | | 8 |
| MP20-25 | 150 | 9 | 12 | 12 | | | 12 |
| MP20-26 | 150 | 9 | 8 | 8 | | | 8 |
| MP20-27 | 150 | 9 | 8 | 12 | 12 | | 12 |
| MP20-28 | 150 | 9 | 12 | 8 | 16 | 8 | 8 |
| MP20-29 | 200 | 9 | 8 | 8 | | | 8 |
| MP20-30 | 200 | 9 | 8 | 12 | 12 | | 12 |

| | | | | | | | |
|---|---|---|---|---|---|---|---|
| MP20-31 | 200 | 9 | 6 | 8 | 12 | 8 | 8 |
| MP20-32 | 200 | 9 | 8 | 6 | 12 | 12 | 12 |
| MP20-33 | 200 | 9 | 12 | 12 | | | 12 |
| MP20-34 | 200 | 10 | 8 | 8 | | | 8 |
| MP20-35 | 250 | 9 | 8 | 8 | | | 8 |
| MP20-36 | 250 | 9 | 12 | 8 | 8 | | 8 |
| MP20-37 | 250 | 9 | 16 | 12 | 12 | | 12 |
| MP20-38 | 250 | 9 | 16 | 12 | 8 | 8 | 8 |
| MP20-39 | 250 | 10 | 16 | 24 | 16 | | 16 |
| MP20-40 | 250 | 10 | 8 | 8 | | | 8 |
| MP20-41 | 250 | 10 | 12 | 12 | | | 12 |
| MP20-42 | 250 | 10 | 16 | 12 | 12 | | 12 |

**Table S13:** Primer sequences used for strain constructions. Underline indicates the first 20 bp of the fluorescent protein gene whereases bold indicates the homology region to the *cat-sacB* cassette.

| Primer name | 5' to 3' | bp |
|---|---|---|
| IS150_PCP25_cs_F | ATAGTACTGTTTAACTTTTCGAGACCTTAGGAGGTAAAAA**CATATGAATATCCTCCTTAGTTCC** | 64 |
| IS150_PCP25_cs_R | TAACCCTTAGTGACTCCTGCAGCGGCCGCTACTAGTATTA **TGTAGGCTGGAGCTGCTTC** | 59 |
| IS150_mtagbfp_F | ATAGTACTGTTTAACTTTTCGAGACCTTAGGAGGTAAAAA <u>ATGAGCGAACTGATCAAAGAGA</u> | 80 |
| IS150_syfp2_F | ATAGTACTGTTTAACTTTTCGAGACCTTAGGAGGTAAAAA <u>ATGGTTAGCAAGGGCGAAGA</u> | 80 |
| IS150_R | CCGAAGACTCTTACTCTTTCAATTTGCAGGCTAAAAACGC ATTACCGCCTTTGAGTGAGC | 80 |
| IS150_screen_F | TAAGATCCCTGCCATTTGG | 19 |
| IS150_screen_R | ATCAAGGAGAAGAAACAAACTT | 22 |

**Table S14:** List of bioinformatic programs, including software versions, used for WGS analysis.

| Tool | Version |
|---|---|
| Abricate | 0.8.10 |
| BWA MEM | 0.7.17-r1188 |
| FastTree | 2.1.10 Double precision (No SSE3): |
| FreeBayes | 1.2.0-dirty |
| IQtree | 1.6.9 for Linux 64-bit built Dec 19 2018 |
| Kraken | 1.0 |
| MLST | 2.16.1 |
| MegaHit | 1.2.9 |
| Newick-Utils | 1.6 |
| Nullarbor | 2.0.20181010 |
| Prokka | 1.13.7 |
| Roary | 3.12.0 |
| SAMtools | 1.9 |
| SKESA | 2.3.0 |
| SPAdes | 3.13.0 |
| Snippy | 4.3.6 and 4.6.0 |
| Trimmomatic | 0.38 |
| centrifuge | 1.0.4 |
| seqret | 6.6.0.0 |
| seqtk | 1.3-r106 |
| snp-dists | 0.6.3 |
| Unicycler | v0.4.8 |
| makeblastdb | 2.9.0+ |
| tblastn | 2.9.0+ |
| bowtie2-build | 2.3.5 |
| bowtie2 | 2.3.5 |
| samtools | 1.9 |
| java | 11.0.1-internal |
| pilon | 1.23 |
| fastp | 0.19.7 |
| fastqc | 0.11.9 |

**Table S15:** Raw data from $IC_{90}$ measurements, presented as modal values throughout the paper. The output values of replicates that met quality control standards were read out on a two-fold scale. Each biological replicate is represented by B and the replicate number.

| Strain | MTX MIC [mg/mL] | | | | | | TMP MIC [µg/mL] | | | | | |
|---|---|---|---|---|---|---|---|---|---|---|---|---|
| | B1 | B2 | B3 | B4 | B5 | Modal | B1 | B2 | B3 | B4 | B5 | Modal |
| DA4201/DA5438 | 16 | 16 | 8 | 32 | | 16 | 0.125 | 0.125 | 0.125 | 0.125 | | 0.125 |
| DA56507 | 16 | 8 | 16 | 8 | 16 | 16 | | | | | | 0.25[1] |
| DA52848 | 16 | 16 | 8 | 8 | 8 | 8 | 0.25 | 0.125 | 0.125 | 0.125 | | 0.125 |
| MP05-31 | >32 | >32 | >32 | | | >32 | >64 | >64 | >64 | | | >64 |
| MP06-01 | | | | | | 4[2] | | | | | | |
| MP06-05 | >32 | >32 | >32 | | | >32 | | | | | | |
| MP06-06 | 32 | 16 | 32 | | | 32 | | | | | | |
| MP06-11 | 4 | 4 | 4 | | | 4 | | | | | | |
| MP06-16 | 16 | 16 | 4 | | | 16 | | | | | | |
| MP06-21 | 4 | 4 | 4 | | | 4 | | | | | | |
| MP06-26 | 16 | 8 | 8 | | | 8 | | | | | | |
| MP06-31 | 16 | 32 | 32 | | | 32 | | | | | | |
| MP06-36 | 4 | 2 | 4 | | | 4 | | | | | | |
| MP06-41 | 8 | 4 | 4 | | | 4 | | | | | | |
| MP06-46 | 32 | 32 | 8 | | | 32 | | | | | | |
| MP18-03 | >32 | >32 | >32 | >32 | | >32 | 4 | 4 | 8 | | | 4 |
| MP18-04 | >32 | >32 | >32 | | | >32 | 8 | 32 | 32 | 32 | 32 | 32 |
| MP18-05 | >32 | >32 | >32 | >32 | | >32 | >64 | >64 | >64 | >64 | >64 | >64 |
| MP18-06 | >32 | >32 | >32 | | | >32 | 8 | 8 | 8 | | | 8 |
| MP18-07 | >32 | >32 | >32 | | | >32 | 32 | 32 | 32 | | | 32 |
| MP18-08 | >32 | >32 | >32 | | | >32 | >64 | >64 | >64 | >64 | >64 | >64 |
| MP18-09 | <2 | <2 | <2 | | | <2 | 0.0625 | 0.0625 | 0.0625 | 0.0625 | | 0.0625 |
| MP18-10 | <2 | <2 | <2 | | | <2 | 0.0625 | 0.0625 | 0.125 | | | 0.0625 |
| MP18-11 | <2 | <2 | <2 | | | <2 | 0.0625 | 0.0625 | 0.0625 | | | 0.0625 |
| MP18-12 | >32 | >32 | >32 | | | >32 | >64 | >64 | >64 | | | >64 |
| MP18-13 | 8 | 8 | 8 | | | 8 | 0.0625 | >0.03125 | >0.03125 | 0.03125 | | >0.03125 |
| MP18-14 | 32 | 32 | 32 | | | 32 | 0.25 | 0.125 | 0.125 | | | 0.125 |
| MP18-15 | 4 | 4 | 4 | | | 4 | 0.25 | 0.125 | 0.125 | | | 0.125 |
| MP18-16 | >32 | >32 | >32 | | | >32 | 0.25 | 0.125 | 0.125 | | | 0.125 |
| MP18-17 | >32 | >32 | >32 | | | >32 | >64 | >64 | >64 | | | >64 |
| MP18-18 | >32 | >32 | >32 | | | >32 | 2 | 2 | 2 | | | 2 |
| MP18-19 | >32 | >32 | >32 | | | >32 | 4 | 4 | 4 | | | 4 |
| MP18-20 | >32 | >32 | >32 | | | >32 | 4 | 4 | 4 | | | 4 |
| MP18-21 | >32 | >32 | >32 | | | >32 | 4 | 4 | 4 | | | 4 |
| MP18-22 | >32 | >32 | >32 | | | >32 | 4 | 4 | 4 | | | 4 |
| MP18-23 | >32 | >32 | >32 | | | >32 | 8 | 8 | 8 | | | 8 |

| | | | | | | | | | | |
|---|---|---|---|---|---|---|---|---|---|---|
| MP18-24 | >32 | >32 | >32 | | >32 | 4 | 4 | 4 | | 4 |
| MP18-25 | >32 | >32 | >32 | | >32 | 8 | 4 | 4 | | 4 |
| MP18-26 | >32 | >32 | >32 | | >32 | 4 | 4 | 4 | | 4 |
| MP18-27 | >32 | >32 | >32 | | >32 | 4 | 4 | 4 | | 4 |
| MP18-28 | >32 | >32 | >32 | | >32 | 4 | 4 | 4 | | 4 |

[1]The MIC for DA56507 was based on the modal from the following nine replicates: 0.25 - 0.25 - 0.125 - 0.125- 0.0625 - 0.0625 - 0.25 - 0.25 - 0.25

[2]MP06-01 was used as an internal standard on all MXT-plates, the MIC was based on the modal from the following 50 replicates: 4 - 4 - 4 - 8 - 4 - 8 - 4 - 4 - 4 - 4 - 8 - 4 - 4 - 4 - 8 - 2 - 4 - 4 - 4 - 4 - 4 - 4 - 8 - 4 - 8 - 8 - 8 - 8 - 4 - 4 - 2 - 2 - 2 - 2 - 4 - 2 - 8 - 4 - 4 - 4 - 4 - 4 - 8 - 4 - 8 - 8 - 8 - 8 - 4 - 8

**Protocols**

Bacterial strains and growth conditions

Bacterial strains used in this study and their assigned reference numbers are listed in Table S1. Most strain constructs were derived from the *E. coli* K-12 strain MG1655 (DA4201/DA5438), or for cloning purposes from *E. coli* DH5α (MP18-09). Two isogenic pairs of clinical *E. coli* strains, where one strain is $TMP^S$ and the other is $TMP^R$, were used for fitness measurements. In the first pair, $TMP^R$ in *E. coli* K56-2 (MP06-01) is caused by one intragenic point mutation T>A (W30R) in the *folA* gene, and one in its promotor region ($P_{folA}$, C>T 58 base pairs (bp) upstream of the gene) (Podnecky et al., 2018, Kahlmeter, 2000) (MP06-05). In the second pair, *E. coli* K56-75 (MP06-41) harbored the pG06-VIM-1 MDR plasmid. This plasmid encodes, amongst other antibiotic resistance genes, two different *dfrA* genes (*dfrA1* and *dfrA12*) as putative $TMP^R$ determinants as well as carbapenem resistance ($bla_{\mathrm{VIM-1}}$) (Di Luca et al., 2016) (MP05-31). All incubations of liquid cultures were performed with orbital shaking (225 rpm) at 37°C, unless otherwise specified. Cultures were grown in Miller Difco Luria-Bertani (LB) broth (Becton, Dickinson and Co.) or SOC (20 g/L tryptone (Oxoid), 5 g/L yeast extract (Oxoid), 0.5 g/L NaCl (VWR), 0.25 mM KCl (Sigma-Aldrich), 10 mM $MgCl_2$ (Sigma-Aldrich), 4 g/L glucose (Sigma-Aldrich). When required, medium was supplemented with 1.5% (w/v) agar (Sigma-Aldrich) to form solid medium. For assays involving either antibiotics or cytostatics, the bacteria were grown in cation adjusted Mueller-Hinton II Broth (MHIIB, Becton, Dickinson and Co.), supplemented with the relevant drug. For assays done on strains harboring the pBAD30 expression vector, cultures were supplemented with 0.2% (w/v) arabinose (Sigma-Aldrich) for induction. Generalized transduction using the P1*vir* (Ikeda and Tomizawa, 1965) were used to move chromosomal markers between strains. For selection against cells expressing *sacB*, sucrose selection plates (5 g yeast extract (Oxoid), 10 g Tryptone (Oxoid), and 1 mM NaOH per liter (VWR), 5 % (w/v) sucrose (Sigma-Aldrich)) were used (Bertani, 1951). When appropriate, media were supplemented with: 12.5 mg/L chloramphenicol (Sigma-Aldrich), 7.5 mg/L tetracycline (Sigma-Aldrich). For long-term storage, strains and populations were mixed with glycerol at a final concentration of 20 % (v/v) and frozen at −80 °C.

Strain Constructions

A promoter-levansucrase-chloramphenicol resistance-promoter cassette ($P_{CP25}$-*sacB*-*cat*-$P_{J23101}$) was first constructed by amplifying the *sacB*-*cat*-$P_{J23101}$ cassette (GenBank accession number KM018298) by using primers with 40 bp sequence homologous to each end of the *insKJ* and partially *mokA* genes in the IS*150* region on the *E. coli* MG1655 chromosome (Table S13). The construct was introduced onto the chromosome by λ Red recombineering (Datsenko and Wanner, 2000, Yu et al., 2000) in a strain carrying the pSIM5 plasmid (Sharan et al., 2009) with tetracycline as the antibiotic selection marker (pSIM5-*tet*, DA45134). Chloramphenicol resistance was used to select for the inserted construct.

Fluorescent protein encoding: *bfp* (*cat*-PJ23101-*mtagBFP2*, blue; (Gullberg et al., 2014); GenBank accession number KM018299), *yfp* (*cat*-PJ23101-*SYFP2*, yellow; (Gullberg et al., 2014); GenBank accession number KM018300) were PCR amplified from previous strains (Gullberg et al., 2014). Both amplifications were carried out using Phusion High-Fidelity DNA Polymerase (Thermo Scientific). Reaction primers were designed with one of the 40 bp homology to the disrupted IS*150* locus whilst the other retained the $P_{CP25}$ promoter (Table S13). Reaction products were purified using the GeneJet Purification Kit (Thermo Scientific) and introduced onto the chromosome by λ Red recombineering by counter-selection on sucrose agar medium. This resulted in [$P_{CP25}$-*sYFP2*] and [$P_{CP25}$-*mtagBFP2*] constructs.

Dup-In methodology of the IS*150* locus was then carried out on all previous constructs (Näsvall et al., 2017) using the *sacB*-*cat*-$P_{J23101}$ cassette (GenBank accession number KM018298) and chloramphenicol resistance as the antibiotic selection. P1*vir* lysates for both fluorescent markers were prepared and transduced into a common background (DA4201) by generalized transduction and segregation of Dup-Ins. Briefly, transduced colonies were picked from plates first with chloramphenicol resistance to transfer the Dup-In with the IS*150* locus, then single colonies were patched on sucrose plates for loss of the *sacB*-*cat*-$P_{J23101}$ cassette but retaining of the IS*150* locus. For screening of the final strains constructed and generation of templates for Sanger sequencing, DreamTaq PCR Master Mix (Thermo Scientific) was used.

The fluorescently tagged strains were then further engineered to obtain $TMP^R$ derivatives. Two point mutations associated with *folA* (the first a non-synonymous point mutation in the *folA* gene, W30R, and the second a point mutation 58 bp upstream of the *folA* gene within the promotor region, C>T) were carefully introduced onto the chromosome of the strains using a double MAGE cycle with the pORTMAGE-2 plasmid as described by its constructors (Nyerges et al., 2016). The carbapenemase-encoding plasmid pG06-VIM-1 (encoding two *dfrA* genes, *dfrA1* and *dfrA12 (Di Luca et al., 2016)*), was transformed into the fluorescently tagged strains using room temperature electroporation (Tu et al., 2016).

To verify the role of *dfrA* genes in both $TMP^R$ as well as methotrexate resistance ($MTX^R$), both *dfrA1* and *dfrA12* were PCR amplified using Phusion High-Fidelity DNA Polymerase (New England BioLabs® Inc.), purified using QIAquick® PCR Purification Kit (QIAGEN), phosphorylated using T4 Polynucleotide Kinase (Thermo Scientific) and cloned using T4 DNA Ligase (Thermo Scientific) into the pBAD30 vector at the *Sma*I site. Thus, the expression of the *dfrA* genes was under the tight inducible control by the $P_{BAD}$ promotor when in the presence of arabinose (Guzman et al., 1995). The purified ligation reactions were transformed into electrocompetent DH5-α cells with electroporation and clones carrying the vector-born *dfrA* genes isolated.

### Susceptibility testing

Due to the bacteriostatic activities of MTX and a lack of a gold standard for MTX microbiological assays we define the MIC of MTX in this study as the 90% inhibitory level ($IC_{90}$). This allows for a high resolution and has previously been used as a proxy for the MIC (Imamovic et al., 2018). The $IC_{90}$ values for TMP were determined as described previously (Podnecky et al., 2018). The $IC_{90}$ values for MXT, using Methotrexate Teva/Ebetrex 100 mg/mL (Pharmachemie B.V./Ebewe Pharma Ges.m.b.H Nfg.KG), were determined using the same protocol with small changes. Briefly, 96-well plates were incubated at 300 rpm (3 mm stroke) for 18 h at 37°C before the $OD_{600}$ was measured using an Epoch 2 Microplate Spectrophotometer (BioTek Instruments, Inc.)/VersaMax™ ELISA Microplate (Molecular Devices®). Internal controls were included on all plates. Percent inhibition was calculated as previously described (Imamovic and Sommer, 2013). All measurements were done using at least three biological replicates where the MIC was set as the most read (modal) value on a two-fold scale of replicates that met quality control standards (Table S2). For characterization of $TMP^R$ mutants isolated from the MTX sub-MIC evolution, TMP MIC was determined by gradient diffusion strips following the manufacturer's guidelines (Liofilchem). Measurements were done using two to four biological replicates, where the MIC was set as the most read (modal) value (Table S12).

### Growth rate measurements

Growth rates were determined using a BioscreenC MBR reader (Oy Growth Curves Ab, Ltd). A minimum of five independent overnight cultures were diluted to ~5 x $10^6$ CFU/mL in MHIIB. Two 300 µL aliquots of each dilution were transferred into sterile Honeycomb plates (Oy Growth Curves Ab, Ltd). The samples were grown at 37°C with continuous shaking for 18 h and $OD_{600}$ values were measured every 4 min. The growth curves from the Bioscreen measurements were analyzed and growth rate calculations done using the statistical software R (Team, 2018) and the Bioscreen Analysis Tool BAT 2.1 (Thulin, 2018). Calculations were based on $OD_{600}$ values between 0.02 and 0.1, where growth was observed to be exponential. Relative growth rates were calculated by dividing the generation time of the sample strain by the generation time of the parental strain.

#### Competition experiments

Competition experiments were performed using the fluorescently tagged strain pairs, both for *folA* and pG06-VIM-1 mediated $TMP^R$. A susceptible strain tagged with either *yfp* or *bfp* was mixed at 1:1 ratio with the constructed $TMP^R$ strains harboring the disparate fluorescence marker to initiate a head-to-head competition, at different MTX concentrations. Six independent cultures (~5 x $10^9$ CFU/mL) of each strain were used to start 12 competitions, i.e. six biological replicates for each color arrangement in a dye-swap set-up. Every 24 h for three to four days the competing strains were passaged 1:1,000 into fresh medium and the mutant to wild type (WT) ratio measured by counting $10^5$ cells using a fluorescence-activated cell sorter (BD FACS Aria III). For safety reasons, all cultures were washed in fresh drug-free MHIIB in order to remove remains of cytostatic drugs from the cultures before being moved over to the cell sorter for analysis. Cells were gently pelleted at 5000 rcf at 4°C for 5 min before removal of the supernatant containing drug, followed by resuspension of the cells in fresh MHIIB.
Selection coefficients were calculated according to the regression model $s=[\ln R(t)/R(0)]/t$, as previously described (Dykhuizen, 1990), where $R$ is the mutant to WT ratio and t is the time measured in generations of growth (Tables S3-S5). The minimum selective concentration (MSC) is defined as the concentration where the selection coefficient equals zero (where the regression line crosses the x-axis) (Gullberg et al., 2011).
In a similar way, six individual cultures of a susceptible *yfp* strain was competed against the *bfp* resistant strains in 1:1, 1:10, 1:$10^2$, 1:$10^3$ and 1:$10^4$ starting ratios of $TMP^R$:$TMP^S$ strains at concentrations slightly above the estimated MSCs (400 µg/mL for *folA* mutant, 75 µg/mL for p06-VIM-1).
To assess the stability of the pG06-VIM-1 in the presence of MTX, three independent lineages of K56-75 harboring the plasmid (MP05-31) were serially passaged for 50 generations (100-fold dilutions) in 1 mL MHII batch cultures amended with 400 µg/mL MTX. The lineages were then plated on non-selective agar and 100 colonies from each lineage replica plated and reduced susceptibilities towards ampicillin, TMP, streptomycin and spectinomycin were scored. Susceptibility was tested by patching on MHII agar, MHII agar supplemented with 100 µg/mL Ampicillin, MHII agar supplemented with 25 µg/mL TMP and MHII agar supplemented with 40 µg/mL streptomycin as well as 40 µg/mL spectinomycin.

#### Selective plating on high concentrations of MTX

Single MTX resistant mutants of K56-2 (MP06-01) were selected at lethal MTX concentrations. Dense overnight cultures grown in drug-free LB was concentrated 10×, and 100 µL spread on LB agar plates supplemented with 4, 8 and 16 mg/mL MTX. Due to the slow growth of mutants, the plates were incubated between 48 to 96 h before mutants were purified on non-selective plates. Additionally, an overnight culture grown in LB containing MTX at the estimated MIC concentration was concentrated 10× and 100 µL spread on LB agar plates with and without MTX 32 mg/mL. After 48 h incubation, mutants were purified on non-selective plates. The MTX and TMP MICs for all mutants isolated were determined as previously described by $IC_{90}$ testing (Podnecky et al., 2018) and the *folA* gene, its promotor area and the *marR* gene sequenced with Sanger sequencing and analyzed using the CLC Main Workbench (Qiagen) (Table S6).

#### Laboratory evolution at sub-MICs of MTX

To examine the effect of MTX presence on $TMP^R$ evolution, strain K56-2 (MP06-01) was serially passaged in liquid cultures with MTX supplemented at concentration slightly above the estimated MSC. Initially, 10 independent overnight cultures were started from independent colonies on separate agar plates. From the original overnight cultures, ~$10^3$ cells were used to start ten independent lineages in 1 mL MHIIB containing 400 μg/mL MTX (lineages 1-10). Every 12 h for 25 days, the lineages were serially passaged by 1,000-fold dilution in 1 mL batch cultures, allowing for ~500 generations of growth. Every ~50 generations the populations were frozen down at –80°C for downstream analysis. In parallel, three independent control lineages were simultaneously sampled for TMP resistance under the same experimental conditions except for MTX exposure (lineages 11-13). All end-point populations were plated on MHII agar plates containing 32 mg/mL MTX after ~500 generations of growth. From lineages 1-10, 20 colonies were isolated from each lineage and tested for $TMP^R$. Of those, none displayed increased resistance towards TMP compared to the parental strain. To examine possible evolution of cross-resistance towards TMP in the laboratory evolution experiment described above, all 13 frozen populations from every 50 generations were gently thawed on ice and dilution series plated on both MHII agar plates containing TMP 4 μg/mL as well as on MHII agar without any drug, and frequencies of TMP resistant mutants calculated. From each plate where mutants grew, up to 5 colonies were randomly isolated, their susceptibility towards TMP measured, and the *folA* gene and its promotor area sequenced with Sanger sequencing and analyzed using the CLC Main Workbench (Qiagen) (Table S12). No $MTX^R$ or $TMP^R$ colonies were isolated from the lineages grown without drug presence (lineage 11-13). This was done to control for the emergence and selection of spontaneous mutants in a drug-free environment.

#### Whole genome sequencing

To investigate the possibility of additional genetic changes, other than the $TMP^R$ determinants shown to be associated with reduced susceptibility towards MTX, during MTX selection, five isolates from the lethal selection were chosen (MP18-13, MP18-17, MP18-20, MP18-26 and MP18-28) based on their different susceptibility profiles and subjected to whole genome sequencing (WGS). Bacteria were grown overnight and genomic DNA prepared using GenElute$^{TM}$ Bacterial Genomic DNA Kit (Sigma-Aldrich) following the protocol provided by manufacturer with slight adaptions. In brief, 1.5 mL of dense culture ($OD_{600}$: 0.8-1.0) was pelleted by centrifugation at 13,000 rpm, culture medium removed and the pellet resuspended in 200 μL lysozyme solution (100 mg/mL) and incubated for 30 min at 37°C before 20 μL of RNase A solution was added and incubated for 2 min at room temperature. Following, 20 μL of Proteinase K (20 mg/mL) and 200 μL of Lysis solution C were added to the mixture and incubated at 55°C for 10 min after being thoroughly vortexed. To each pre-assembled GenElute Miniprep Binding Column, 500 μL of the Column Preparation Solution were added, 200 μL of ethanol (95-100 %) was then added to the lysate and thoroughly mixed before the lysate was carefully loaded onto the binding column, centrifuged at 13,000 rpm for 1 min and then washed 2× with 500 μL of Wash Solution. Genomic DNA was eluted in 100 μL of 10 mM Tris-base and purity and concentration determined using NanoDrop One (Thermo Scientific) and Qubit (Thermo Scientific) respectively. Next-generation sequencing libraries were prepared from the bacterial genomic DNA samples and sequenced on an Illumina NovaSeq with a 2 × 150 bp configuration (GENEWIZ). Average whole genome coverage per sample was approximately 700. Analysis of the fastq files obtained from Illumnia sequencing was performed using an in-house bioinformatic pipeline (Table S14) to compare the mutant sequences to the previously published WT strain (available at NCBI, BioSample SAMN08095529). Where single-nucleotide polymorphisms (SNPs) were identified with a coverage below 100 the evidence was considered insufficient and the SNPs were removed from the analysis. Raw sequence reads were deposited under BioProject PRJNA677979.

BERTANI, G. 1951. Studies on lysogenesis. I. The mode of phage liberation by lysogenic Escherichia coli. *Journal of bacteriology,* 62**,** 293-300.

DATSENKO, K. A. & WANNER, B. L. 2000. One-step inactivation of chromosomal genes in <em>Escherichia coli</em> K-12 using PCR products. *Proceedings of the National Academy of Sciences,* 97**,** 6640.

DI LUCA, M. C., SAMUELSEN, Ø., STARIKOVA, I., KLOOS, J., HÜLTER, N., JOHNSEN, P. J., SØRUM, V. & NASEER, U. 2016. Low biological cost of carbapenemase-encoding plasmids following transfer from Klebsiella pneumoniae to Escherichia coli. *Journal of Antimicrobial Chemotherapy,* 72**,** 85-89.

DYKHUIZEN, D. E. 1990. Experimental Studies of Natural Selection in Bacteria. *Annual Review of Ecology and Systematics,* 21**,** 373-398.

GULLBERG, E., ALBRECHT, L. M., KARLSSON, C., SANDEGREN, L. & ANDERSSON, D. I. 2014. Selection of a Multidrug Resistance Plasmid by Sublethal Levels of Antibiotics and Heavy Metals. *mBio,* 5**,** e01918-14.

GULLBERG, E., CAO, S., BERG, O. G., ILBÄCK, C., SANDEGREN, L., HUGHES, D. & ANDERSSON, D. I. 2011. Selection of Resistant Bacteria at Very Low Antibiotic Concentrations. *PLOS Pathogens,* 7**,** e1002158.

GUZMAN, L. M., BELIN, D., CARSON, M. J. & BECKWITH, J. 1995. Tight regulation, modulation, and high-level expression by vectors containing the arabinose PBAD promoter. *Journal of bacteriology,* 177**,** 4121-4130.

IKEDA, H. & TOMIZAWA, J.-I. 1965. Transducing fragments in generalized transduction by phage P1: I. Molecular origin of the fragments. *Journal of Molecular Biology,* 14**,** 85-109.

IMAMOVIC, L., ELLABAAN, M. M. H., DANTAS MACHADO, A. M., CITTERIO, L., WULFF, T., MOLIN, S., KROGH JOHANSEN, H. & SOMMER, M. O. A. 2018. Drug-Driven Phenotypic Convergence Supports Rational Treatment Strategies of Chronic Infections. *Cell,* 172**,** 121-134.e14.

IMAMOVIC, L. & SOMMER, M. O. A. 2013. Use of Collateral Sensitivity Networks to Design Drug Cycling Protocols That Avoid Resistance Development. *Science Translational Medicine,* 5**,** 204ra132.

KAHLMETER, G. 2000. The ECO.SENS Project: a prospective, multinational, multicentre epidemiological survey of the prevalence and antimicrobial susceptibility of urinary tract pathogens--interim report. *J Antimicrob Chemother,* 46 Suppl 1**,** 15-22; discussion 63-5.

NÄSVALL, J., KNÖPPEL, A. & ANDERSSON, D. I. 2017. Duplication-Insertion Recombineering: a fast and scar-free method for efficient transfer of multiple mutations in bacteria. *Nucleic Acids Res,* 45**,** e33.

NYERGES, Á., CSÖRGŐ, B., NAGY, I., BÁLINT, B., BIHARI, P., LÁZÁR, V., APJOK, G., UMENHOFFER, K., BOGOS, B., PÓSFAI, G. & PÁL, C. 2016. A highly precise and portable genome engineering method allows comparison of mutational effects across bacterial species. *Proceedings of the National Academy of Sciences,* 113**,** 2502.

PODNECKY, N. L., FREDHEIM, E. G. A., KLOOS, J., SØRUM, V., PRIMICERIO, R., ROBERTS, A. P., ROZEN, D. E., SAMUELSEN, Ø. & JOHNSEN, P. J. 2018. Conserved collateral antibiotic susceptibility networks in diverse clinical strains of Escherichia coli. *Nature Communications,* 9**,** 3673.

SHARAN, S. K., THOMASON, L. C., KUZNETSOV, S. G. & COURT, D. L. 2009. Recombineering: a homologous recombination-based method of genetic engineering. *Nature protocols,* 4**,** 206-223.

TEAM, R. C. 2018. R: A language and environment for statistical computing. R Foundation for Statistical Computing, Vienna, Austria.

THULIN, M. 2018. *BAT: an online tool for analysing growth curves.* [Online]. Available: http://www.mansthulin.se/bat/ [Accessed 2019].

TU, Q., YIN, J., FU, J., HERRMANN, J., LI, Y., YIN, Y., STEWART, A. F., MÜLLER, R. & ZHANG, Y. 2016. Room temperature electrocompetent bacterial cells improve DNA transformation and recombineering efficiency. *Scientific reports,* 6**,** 24648-24648.

YU, D., ELLIS, H. M., LEE, E. C., JENKINS, N. A., COPELAND, N. G. & COURT, D. L. 2000. An efficient recombination system for chromosome engineering in Escherichia coli. *Proceedings of the National Academy of Sciences of the United States of America,* 97**,** 5978-5983.
